# Cell-type specific molecular signatures of aging revealed in a brain-wide transcriptomic cell-type atlas

**DOI:** 10.1101/2023.07.26.550355

**Authors:** Kelly Jin, Zizhen Yao, Cindy T. J. van Velthoven, Eitan S. Kaplan, Katie Glattfelder, Samuel T. Barlow, Gabriella Boyer, Daniel Carey, Tamara Casper, Anish Bhaswanth Chakka, Rushil Chakrabarty, Michael Clark, Max Departee, Marie Desierto, Amanda Gary, Jessica Gloe, Jeff Goldy, Nathan Guilford, Junitta Guzman, Daniel Hirschstein, Changkyu Lee, Elizabeth Liang, Trangthanh Pham, Melissa Reding, Kara Ronellenfitch, Augustin Ruiz, Josh Sevigny, Nadiya Shapovalova, Lyudmila Shulga, Josef Sulc, Amy Torkelson, Herman Tung, Boaz Levi, Susan M. Sunkin, Nick Dee, Luke Esposito, Kimberly Smith, Bosiljka Tasic, Hongkui Zeng

## Abstract

Biological aging can be defined as a gradual loss of homeostasis across various aspects of molecular and cellular function. Aging is a complex and dynamic process which influences distinct cell types in a myriad of ways. The cellular architecture of the mammalian brain is heterogeneous and diverse, making it challenging to identify precise areas and cell types of the brain that are more susceptible to aging than others. Here, we present a high-resolution single-cell RNA sequencing dataset containing ∼1.2 million high-quality single-cell transcriptomic profiles of brain cells from young adult and aged mice across both sexes, including areas spanning the forebrain, midbrain, and hindbrain. We find age-associated gene expression signatures across nearly all 130+ neuronal and non-neuronal cell subclasses we identified. We detect the greatest gene expression changes in non-neuronal cell types, suggesting that different cell types in the brain vary in their susceptibility to aging. We identify specific, age-enriched clusters within specific glial, vascular, and immune cell types from both cortical and subcortical regions of the brain, and specific gene expression changes associated with cell senescence, inflammation, decrease in new myelination, and decreased vasculature integrity. We also identify genes with expression changes across multiple cell subclasses, pointing to certain mechanisms of aging that may occur across wide regions or broad cell types of the brain. Finally, we discover the greatest gene expression changes in cell types localized to the third ventricle of the hypothalamus, including tanycytes, ependymal cells, and *Tbx3*+ neurons found in the arcuate nucleus that are part of the neuronal circuits regulating food intake and energy homeostasis. These findings suggest that the area surrounding the third ventricle in the hypothalamus may be a hub for aging in the mouse brain. Overall, we reveal a dynamic landscape of cell-type-specific transcriptomic changes in the brain associated with normal aging that will serve as a foundation for the investigation of functional changes in the aging process and the interaction of aging and diseases.

## Introduction

Mammalian brains can display remarkable stability and vulnerability to aging-related decline. Various aspects of behaviors remain robust as animals age, while other functions exhibit marked age-associated decline. The decline in proficiency and performance, including many motor and cognitive tasks, can be dramatically exacerbated by neurodegenerative diseases^1^. Furthermore, age is the major risk factor for these neurodegenerative diseases, such as Alzheimer’s disease and Parkinson’s disease^1^.

Defining and distinguishing global, region-specific, as well as cell-type specific functional changes with age is an essential step towards understanding both the normal aging process and the interaction between normal aging and pathology. In the past decade, there have been concerted efforts to document and catalogue various molecular and cellular hallmarks of aging that are conserved across different model systems^2,3^. Indeed, emerging studies of brain aging and neurodegeneration are beginning to reveal the presence of some of these hallmarks of aging across the brain, including chronic inflammation mediated by microglia and other glial types in the brain^4,5^, cellular senescence^6^, and others^3^. While these hallmarks provide a crucial foundational understanding of how individual cells age, our understanding of how a multicellular tissue as complex and heterogeneous as the brain ages is still rudimentary. We have barely begun to uncover the cellular hallmarks of aging at the cell-type level, and how these changes ultimately contribute to the decline in health of the entire organism.

To address these challenges, many have turned toward single-cell resolution sequencing approaches. In recent years, several studies profiled transcriptomic changes during normal aging across the broad regions of the mouse brain at single-cell level^7,8^, and many more studies profiled more targeted, specific regions or cell types^4,9–15^. While these studies varied in approach and scale, they consistently demonstrated heterogeneity in transcriptomic changes that different cell types display with age. As such, detailed annotation and interrogation of all cell types in the brain will be crucial to fully characterize how different cell types, both neuronal and non-neuronal, change and interact with one another during aging.

Despite tremendous advances in single-cell brain aging research, many challenges remain. Studies on the whole brain or very large portions of the brain often lacked cell type resolution and sequencing depth to cover diverse cell types. On the other hand, studies targeting smaller brain regions were usually conducted by different groups under variable conditions, making it difficult to compare and integrate the studies into a consistent view. Most recently, scaling single-cell transcriptomic approaches to the whole mouse brain has allowed us to define cell types in the brain at an unprecedented resolution and comprehensiveness, revealing the tremendous diversity of neuronal and non-neuronal cell types and their gene expression profiles throughout the adult mouse brain^16–19^. These studies present a timely opportunity to obtain a systematic and comprehensive understanding of how the brain changes with age at molecular and cellular levels.

Here, we use single-cell RNA sequencing (scRNA-seq) to profile a wide range of brain regions covering major parts of the brain that have complex cell type compositions, in young adult (2 months old) and aged (18 months old) mice in both sexes. Together, these profiled regions cover approximately 35% of the entire volume of the mouse brain. The total dataset includes ∼1.2 million high-quality single-cell transcriptomes from young adult and aged mice that have been annotated using the Allen whole mouse brain cell type atlas (companion paper Yao *et. al.*^17^), allowing us to identify over 130 unique transcriptomic subclasses (which can be further subdivided into many more supertypes and clusters) and interrogate them for age-associated gene expression changes. We also present two spatial transcriptomics datasets that focus on specific cell types in specific regions of interest.

In this study, we confirm and extend upon previous studies observing greatest gene expression changes with age in many non-neuronal types. In addition, we discover changes in types that have not been majorly implicated in brain aging in the past. In particular, we find a large number of age-associated gene expression changes in both neuronal and non-neuronal types surrounding the third ventricle of the hypothalamus, including tanycytes, ependymal cells, and neurons in the arcuate nucleus (ARH). Many of the cell types with the greatest gene expression changes are known for their roles in nutrient and energy homeostasis, including neuronal types that express *Agrp* and *Pomc*, markers of neurons involved in the central melanocortin signaling circuit. Taken together, our results systematically reveal a wide range of cell-type specific patterns of aging, identify age-specific cell type clusters that show unique gene expression changes, and highlight the third ventricle area of the hypothalamus as a potential hot spot for brain aging, likely via its role in dysregulation of nutrient sensing and homeostasis, one of the known hallmarks of aging^2^.

## Results

### Brain-wide single-cell and *in situ* RNA profiling in aged and adult mouse brain

To evaluate cell-type specific transcriptomic changes with age, we profiled 16 broadly dissected regions across the young adult (P56; 2-month-old) and aged (P540; 18-month-old) female and male mouse brains using 10x Genomics Chromium platform based on version 3 chemistry (10xv3). These 16 broad regions (**Figure 1a**) were selected due to their known sensitivity to age and age-associated diseases in the literature^20^. They were grouped into six major brain structures: 1) isocortex, which includes prelimbic area + infralimbic area + orbital area (PL + ILA + ORB), agranular insular area (AI), anterior cingulate area (ACA), and retrosplenial area (RSP); 2) hippocampal formation (HPF), which includes hippocampus (HIP), parasubiculum + postsubiculum + presubiculum + prosubiculum + subiculum (PAR + POST + PRE + ProS + SUB), and lateral and medial entorhinal areas (ENT); 3) hypothalamus (HY); 4) cerebral nuclei (CNU), which includes the dorsal and ventral striatum (STRd, STRv), pallidum (PAL), and striatum-like amygdalar nuclei (sAMY); 5) midbrain, which includes periaqueductal gray + midbrain raphe nuclei (PAG + RAmb) as well as substantia nigra + ventral tegmental area (SNr + SNc + VTA); 6) hindbrain, which includes the anterior or posterior part of the combined pons, motor related and behavioral state related areas (Pmot/sat–A; Pmot/sat-P). Brain regions for profiling and boundaries for dissections were defined by Allen Mouse Brain Common Coordinate Framework version 3 (CCFv3)^21^ as previously described^16^ (**Figure 1a,b**). Based on three-dimensional volumes as estimated by CCFv3, we estimate that these 16 broad dissection regions, encompassing ∼110 CCF-defined brain regions, cover approximately 35% of all grey matter areas within the whole mouse brain.

**Figure 1.**
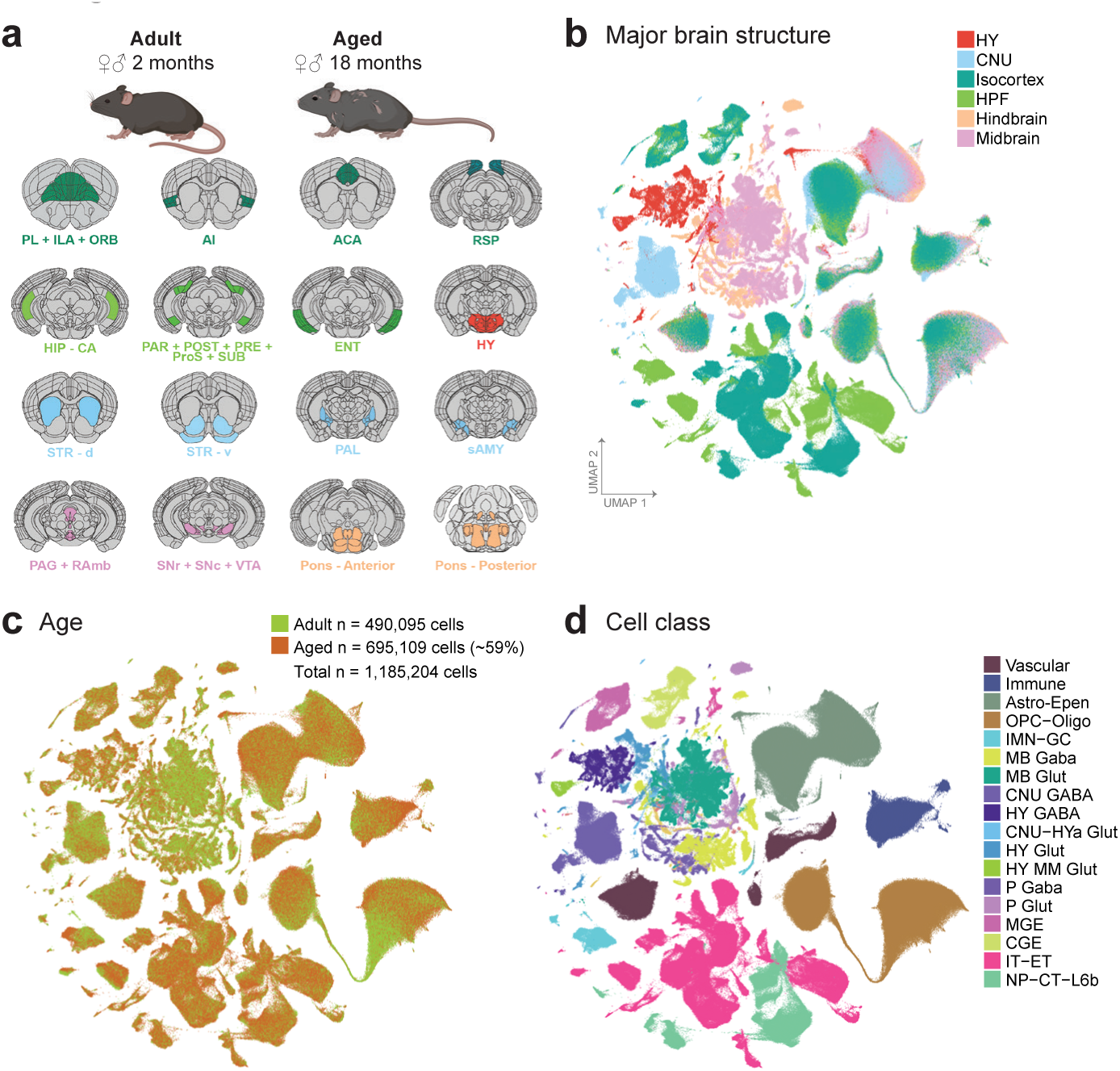
Transcriptomic cell types in the aged and adult mouse brain. **(a)** Schematic of dissected brain regions profiled in this study, colored by major brain structure. **(b-c)** UMAP representation of n = 1,185,204 cells included in this study, colored by major brain structure (b) and cell class (c). Mouse depictions in (a) are created with BioRender.com.

Our final dataset includes single-cell transcriptomes from 272 unique 10xv3 libraries, which were collected from a total of 96 mice (**Supplementary Table 1**). To ensure good representation of both neurons and non-neuronal cells, we employed multiple forms of fluorescence-activated cell sorting (FACS) and unbiased cell sampling (labeled as “No FACS”; **Methods**). All neuron-enriched libraries were FACS-isolated from the pan-neuronal *Snap25-IRES2-Cre/wt;Ai14/wt* transgenic mice, whereas the unbiased libraries were isolated from a mixture of transgene-positive and negative mice (**Supplementary Table 1**).

Low-quality transcriptomes were removed based on a combination of quality control (QC) criteria (e.g., gene detection, qc score, and doublet score, see **Methods; Extended Data Figure 1a**). After the QC-filtering, we obtained 1,185,204 high-quality cells, of which ∼59% (695,109 cells) originated from aged, and the rest (490,095 cells) from young adult brain tissue (**Extended Data Figure 1a**). Post QC-filtering, we assessed a variety of quality scores, including gene detection, QC score, and mitochondrial RNA percentage (mito score) and observed little variation between aged and adult cells for most cell classes (**Extended Data Figure 1b-d**), giving us confidence that tissue age did not significantly affect the quality of sequencing libraries. We only observed differences in these metrics for a small number of cell classes, such as higher gene detection in adult IMN-GC (immature neurons and granule cells) compared to aged IMN-GC (**Extended Data Figure 1b)**.

Following QC, we performed *de novo* clustering of all adult and aged cells together (**Methods; Extended Data Figure 1a**). Briefly, all the adult cells in this study had been thoroughly annotated as part of our recent mouse whole brain taxonomy^17^, allowing us to leverage the existing cell type annotations to help annotate the aged cells. Aged cells that co-clustered with an adult cell type that made up greater than 10% of the cluster were assigned the majority identity from the adult cells at the subclass level. All cells in this study have at least 3 levels of annotation: 1) cell category (the broadest level of annotation), 2) class, and 3) subclass. The subsequent figures of this study will highlight certain populations of cells for which additional clustering was performed and finer-level cell type annotations were assigned including 4) supertype, and 5) cluster, which is the finest level of annotation we use.

Out of the total 306 subclasses defined in our whole mouse brain cell atlas^17^, we identified a total of 185 unique subclasses in the combined aged and adult dataset. Of those 185 subclasses, 132 subclasses met our criteria to include in downstream analysis for age differential gene expression (**Methods**). These 132 subclasses spanned 18 different cell classes (**Figure 1c; Supplementary Table 2**) and displayed specific marker gene expression (**Extended Data Figure 2**). Slightly more than half of all cells in this study were non-neuronal, and their proportion varied by brain region (**Supplementary Table 2; Extended Data Figure 1e**). Most non-neuronal cell types were shared between brain regions, whereas neurons differed among brain regions (**Figure 1b,c**; **Figure 2**). We also observed that not all subclasses were perfectly balanced between ages and sexes, as is expected for this type of data (**Figure 1b**, **Figure 2; Supplementary Table 2**). The ratios of age and sex for each subclass are summarized in **Figure 2** and **Supplementary Table 2**.

**Figure 2.**
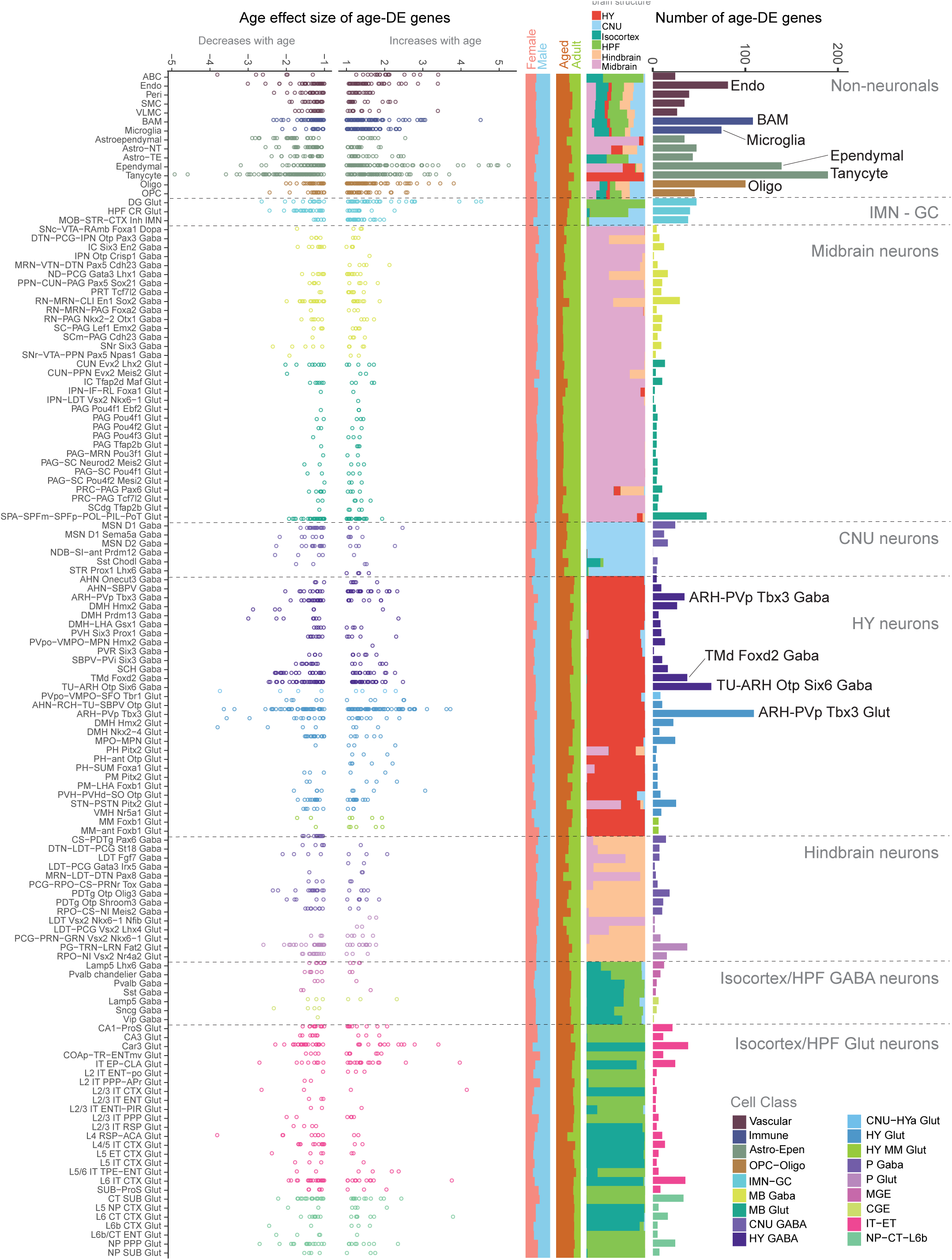
Differentially expressed genes across cell subclasses in the aged and adult mouse brain. Summary of the number and effect size of all age-DE genes identified at the subclass level. Far right: The total number of age-DE genes within each subclass, colored by cell class and ordered based on broad categories. Center: Bar charts that summarize the breakdown of each subclass by major brain structure, age, and sex. Far left: Age effect sizes of all age-DE genes for each subclass.

To complement the scRNA-seq data, we collected two separate Molecular Cartography datasets (a form of *in situ* spatial RNA profiling from Resolve Biosciences) to visualize and validate results discovered by scRNA-seq. For each spatial dataset, we selected a panel of 100 genes to profile pre-selected region(s) in male and female mouse coronal brain sections. These two datasets span a variety of different areas including regions in the isocortex, striatum, hindbrain, midbrain, and hypothalamus, and will be referred to in the remainder of the text as Resolve spatial transcriptomics experiments 1 and 2 (RSTE1,2 in **Extended Data Figure 3a,b**).

### Analysis of age-associated differential gene expression across subclasses

To examine and model age-associated differentially expressed genes (age-DE genes) within each subclass, we used Model-based Analysis of Single-cell Transcriptomics (MAST^22^) with two different statistical models as described in **Methods**. Briefly, due to the variability of FACS population plans and genotypes across aged and adult libraries (**Extended Data Figure 4a**), and the fact that cells from different FACS population plans were observed to have an effect on quality metrics such as gene detection and QC score (**Extended Data Figure 4b,c**), we used two different statistical models with different covariates to try to account for these differences (**Methods**). Age effect size, which can be interpreted as an estimate of log_2_ fold change with age, and adjusted p-value were calculated from the model. Age effect sizes as estimated by these two models were found to vary for certain subclasses, with neuronal subclasses showing a greater variation than non-neuronal ones, likely due to the smaller number of libraries contributing to each neuronal subclass (**Extended Data Figure 4d; Supplementary Table 3**). As a result, we implemented a stringent set of significance criteria – only genes found to be significant with an |age effect size| > 1 and p-value < 0.01 under both models were considered significant and reported here. Positive age effect sizes (> 1) roughly correspond to an increase of more than two-fold in that gene with age, while negative age effect sizes (< −1) roughly correspond to a decrease of more than 50%. Age effect sizes and p-values from both models for each significant gene are reported in **Supplementary Table 3.**

Across the 132 subclasses included in this analysis, we found over 1,200 unique age-DE genes, many of which in non-neuronal subclasses, and comparatively fewer within most neuronal subclasses (**Figure 2; Supplementary Tables 2,3**). Within the non-neuronal subclasses, the greatest numbers of age-DE genes were found in tanycytes and ependymal cells, which both belong to the Astro-Epen cell class. Across the neuronal subclasses, the greatest numbers of age-DE genes were found in hypothalamic subclasses (**Figure 2; Supplementary Tables 2,3**).

Across all subclasses, we found that the vast majority of age-DE genes were significant in only one or two subclasses (**Extended Data Figure 5a**), suggesting that most age-DE genes were cell type specific. We also found a handful of age-DE genes with significant changes in many subclasses (**Extended Data Figure 5a**), and many of these genes displayed region and/or cell-type specific differential expression. For example, *3222401L13Rik* (a long intergenic non-coding RNA^23^ surrounded by protocadherins in the genome) and *Slc5a5* (a gene encoding a sodium/iodide cotransporter) were significantly upregulated in 70 and 48 subclasses, respectively, almost all of which were midbrain, hindbrain, and hypothalamic neuronal types (**Extended Data Figure 5b**). We also observed increased expression of *AC149090.1* in an even wider array of regions and types (54 subclasses), including cortical neurons and glial types (**Extended Data Figure 5b**). *AC149090.1* is an ortholog of *Pisd* which encodes phosphatidylserine decarboxylase, an enzyme involved in lipid metabolism^24^ linked to mitochondrial disease^25^. *AC149090.1* was also the top contributing gene in a recent study that built cell-type specific transcriptomic age clocks from scRNA seq data in mouse subventricular zone^14^. We also observed genes that decreased with age across multiple subclasses, including *Ccnd1* and *Ccnd2* that encode cell cycle regulator proteins cyclin D1 and D2 respectively, decreasing with age in various hypothalamic neuronal subclasses, particularly ones localized to the periventricular area of the hypothalamus including the dorsomedial nucleus (DMH) and ARH (**Extended Data Figure 5b**). Altogether, these observations suggest that different subclasses demonstrate unique combinations of gene expression profiles that are influenced by age.

### Changes in OPCs and Oligodendrocytes with age

Mature oligodendrocytes are the myelinating cells of the brain. They make up most of the white matter in the brain by creating and maintaining the myelin sheaths that encase and protect axons within the central nervous system. Oligodendrocytes develop from oligodendrocyte precursor cells (OPCs). Brain-wide decrease in white matter volume with normal aging has been well-characterized^26,27^ and correlates with cognitive decline^28,29^.

We profiled 88,535 OPCs and 165,858 oligodendrocytes in our scRNA-seq dataset. To obtain cell identities at the finer supertype level, we mapped our oligodendrocyte population to an scRNA-seq dataset generated by Marques *et al.*^30^. We resolved our oligodendrocyte population into the following supertypes: committed oligodendrocyte precursors (COP), newly formed oligodendrocytes (NFOL), myelin-forming oligodendrocytes (MFOL), and mature oligodendrocytes (MOL). We saw a smooth transition from OPC to MOL in the UMAP space (**Figure 3a**), as well as separation of cells by age and region. Separation by age was most striking within the MOL cell population, whereas the separation by region was more apparent in OPCs (**Figure 3a**).

**Figure 3.**
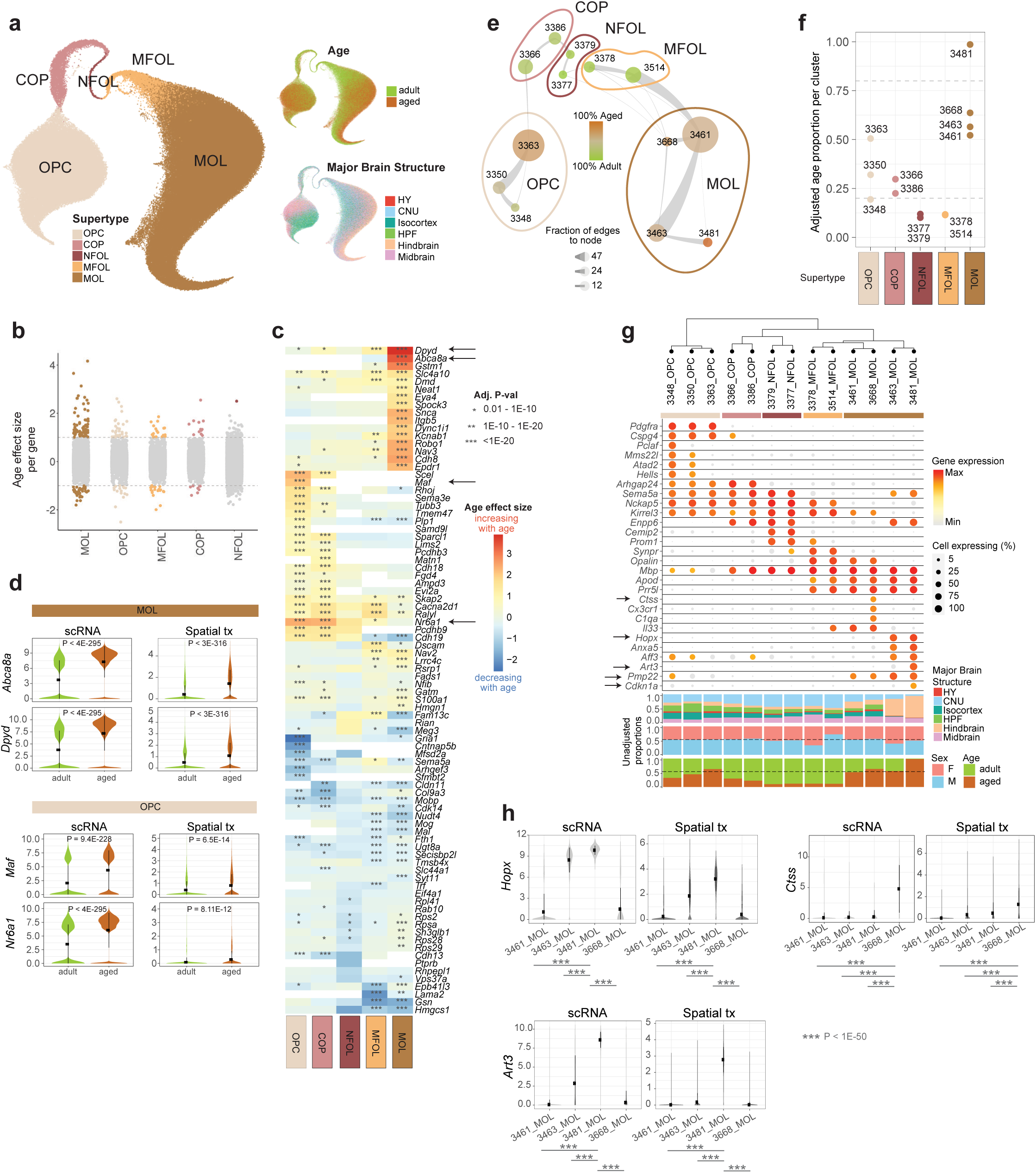
Age-associated changes in OPCs and oligodendrocytes. **(a)** UMAP of all OPC and oligodendrocyte transcriptomes colored by supertype, age, and major brain structure. **(b)** Age effect sizes of age-DE genes within OPCs and oligodendrocyte supertypes, with significant age-DE genes colored (absolute age effect size >1 and P < 0.01). **(c)** Heatmap of age effect sizes of top age-DE genes within OPCs and oligodendrocyte supertypes. Asterisks denote statistical significance. **(d)** Violin plots of expression of *Abca8a* and *Dpyd* in MOL and *Maf* and *Nr6a1* in OPC from scRNA-seq and spatial RSTE1 datasets. **(e)** Constellation plot representing OPC and oligodendrocyte clusters using UMAP coordinates shown in (a). Node (cluster) size is proportional to cell number. Edge thickness is proportional to the fraction of nearest neighbors that were assigned to the connecting node scaled to node size. Cluster color represents the percent of aged or adult cells. **(f)** Adjusted age proportion of each cluster from (e), colored and grouped by supertype. **(g)** Dendrogram and dot plot of cluster marker genes. Below dot plot are bar summaries of each cluster broken down by major brain structure, sex, and age. Dendrogram is calculated from cluster DE genes. **(h)** Violin plot expression of *Hopx*, *Art3,* and *Ctss* in MOL clusters from scRNA-seq and spatial dataset RSTE1.

We found the greatest number of age-DE genes in MOL, followed by OPC, and then MFOL (**Figure 3b**). The signatures of age-DE genes between OPC and COP resembled each other, while those between MFOL and MOL most resembled each other. This is consistent with their developmental trajectory and relatedness to one another in the UMAP space (**Figure 3a,c**). Amongst these age-DE genes, there was a strong increase in expression of *Abca8a* and *Dpyd* across MOL (**Figure 3c**), which was confirmed with spatial transcriptomics dataset RSTE1 (**Figure 3d**). *Abca8a* is the mouse homolog of human ABCA8, a gene known for its ability to stimulate sphingomyelin production and regulate lipid metabolism in oligodendrocytes in humans^31^. *Dpyd* encodes an enzyme involved in the breakdown of pyrimidines, and has also been shown to play a role in lipid degradation^32^. Increase in expression of both genes with age points to an alteration in myelin maintenance capacity in MOL with age. We also observed and spatially confirmed the increased expression of *Maf* and *Nr6a1* in OPC (**Figure 3c,d**). *Maf* encodes a transcription factor that heterodimerizes with transcription factor *Nrf2*, a master regulator of redox status, antioxidative, and anti-inflammatory response^33^. Altered levels of *Nrf2* and *Maf* expression in the brain have been associated with cognitive impairment and OPC senescence^33,34^.

We tested whether any gene ontology (GO) terms were enriched in genes that were significantly up- or down regulated across different supertypes. We found an enrichment in ion channel activity in downregulated age-DE genes in OPCs, while genes involved in transporter activity and metal ion transport were upregulated in MFOL with age (**Extended Data Figure 6a; Supplementary Table 4**). In MOL, we observed an enrichment of GO terms related to locomotory behavior and neuronal structure-related terms such as synaptic cleft and dendrite development in genes upregulated with age, as well as enrichment of GO terms related to myelin sheath in genes that decreased with age, suggesting that myelin sheath integrity may be compromised with age (**Extended Data Figure 6a; Supplementary Table 4**), a pattern that has also been observed in the transcriptomes of human Alzheimer’s disease brain cells^35^.

We further clustered the data to explore finer (cluster-level) cell types within OPCs and oligodendrocytes. This resulted in 13 transcriptionally distinctive clusters, 3 of which were OPCs, 4 that were MOLs, and the remaining 6 from the transitioning supertypes (**Figure 3e**). To assess whether any of the clusters were age-(>80% adjusted age proportion) or adult-biased (<20% adjusted age proportion), we calculated the adjusted age proportion of each cluster by normalizing to the subclass-wide age proportion (**Methods**). We observed that all transitioning supertypes (COP, NFOL, MFOL) were composed of fewer than 30% aged cells, with NFOL and MFOL clusters being more adult-biased than COP (**Figure 3e,f**). This is consistent with the reported decrease in OPC differentiation with age^36,37^. To confirm these changes in abundance of oligodendrocyte supertypes in the brain with age *in situ*, we calculated the proportion of each supertype in spatial transcriptomics dataset RSTE1 from cortex, striatum, midbrain, and hindbrain (**Extended Data Figure 6b**). We found that while there was no significant change in OPC proportions across regions with age, there was a significant decrease in the proportions of cells in transitionary oligodendrocyte supertypes (COP, NFOL, and MFOL) with age (**Extended Data Figure 6b**), consistent with age proportions observed in scRNA-seq oligodendrocyte clusters (**Figure 3e,f**). In contrast, we observed significant increase in MOL proportions across all imaged brain regions with age in the spatial data (**Extended Data Figure 6b**) as well as the MOL proportions calculated from unbiased scRNA seq libraries (**Extended Data Figure 6c**), consistent with observations of increased MOL accumulation with age made by others^38,39^.

Upon examining marker genes for clusters, we observed expected expression of canonical OPC marker genes such as *Cspg4* (NG2 in humans) across all OPC clusters, *Apod* and *Prr5l* across MFOL and MOL clusters, and increasing *Mbp* expression as OPCs develop on their path to maturity (**Figure 3g**). Across the 3 OPC clusters, we found a graded decrease in DNA repair/chromatin binding genes such as *Hells*, *Atad2*, and *Mms22l* that correlated with the age proportion of each cluster. In MOL, we found two clusters, 3463 and 3481, that were both enriched for hindbrain cells, consistent with increased expression of *Pmp22*, a peripheral myelin gene, high levels of which are typically associated with the myelinating Schwann cells of the peripheral nervous system, and at relatively lower levels in the hindbrain and spinal cord^40^ (**Figure 3g**). Unexpectedly, these hindbrain MOL clusters do not express *Opalin*, a gene commonly considered as a MFOL and MOL-specific marker^41,42^ (**Figure 3g**). Furthermore, both clusters express unique markers that are absent from other MOL clusters, including *Hopx* and *Anxa5*. One of these MOL clusters, 3481, is an age-biased cluster (**Figure 3f**) and expresses a unique gene marker, *Art3*. We confirmed this age-related enrichment of *Art3* by spatial transcriptomics (**Figure 3h**). This observation suggests that MOLs from the hindbrain regions may age differently from MOLs in other brain areas. Also of note, cluster 3481 shows high expression of cell cycle gene *Cdkn1a* (**Figure 3g**), also known as p21, whose increased expression is often associated with cellular senescence^3,43^. While senescent astrocytes and microglia have been observed in the aging brain, whether or not oligodendrocytes undergo cellular senescence in the aged brain remains unclear^6^. As such, cluster 3481 may be a novel, previously uncharacterized type of MOL related to senescence. We also observed a MOL cluster (3668) that is enriched for canonical microglia markers including *Cx3cr1*, *Ctss*, and *C1qa* (**Figure 3g**), possibly representing a cluster of cells with increased inflammation signals and recruitment of microglia. This cluster was detected in spatial dataset RSTE1 across all 4 profiled regions. The proportion of this cluster within the MOL supertype increased with age (**Extended Data Figure 6c**) as well as expression of microglia marker *Ctss* compared to other MOL clusters (**Figure 3h**). Altogether, this analysis confirms previously observed decrease in MOL development with age, as well as identifies, to our knowledge, two novel *Opalin*-negative MOL clusters that are enriched in the hindbrain, one of which is specifically enriched in aged hindbrain and displays markers of cellular senescence.

### Changes in microglia and macrophages with age

In our scRNA-seq dataset, we annotated microglia, border-associated macrophages (BAM), lymphoid cells, and dendritic cells, all belonging to the Immune cell class (**Figure 4a**). Due to limited numbers of lymphoid and dendritic cells, we focused the analysis of immune cells on microglia and BAM. Although we detected far fewer BAMs (n = 3,109 cells) than microglia (n = 69,258 cells) in the scRNA-seq dataset, we observed a greater number of age-DE genes in BAMs than microglia (**Figure 2**). At the subclass level, BAMs showed coordinated upregulation of many *Cd209* genes, which code for lectins that function in cell adhesion and pathogen recognition (**Figure 4b**). From GO analysis, we found upregulated terms with age, enriched in Cd209 genes including carbohydrate binding, lymphocyte proliferation, virus receptor activity, and others (**Figure 4d**, **Supplementary Table 4**). An increase in *Cd209a* and *Cd209b* with age was confirmed by spatial transcriptomics (dataset RSTE1, **Figure 4c**).

**Figure 4.**
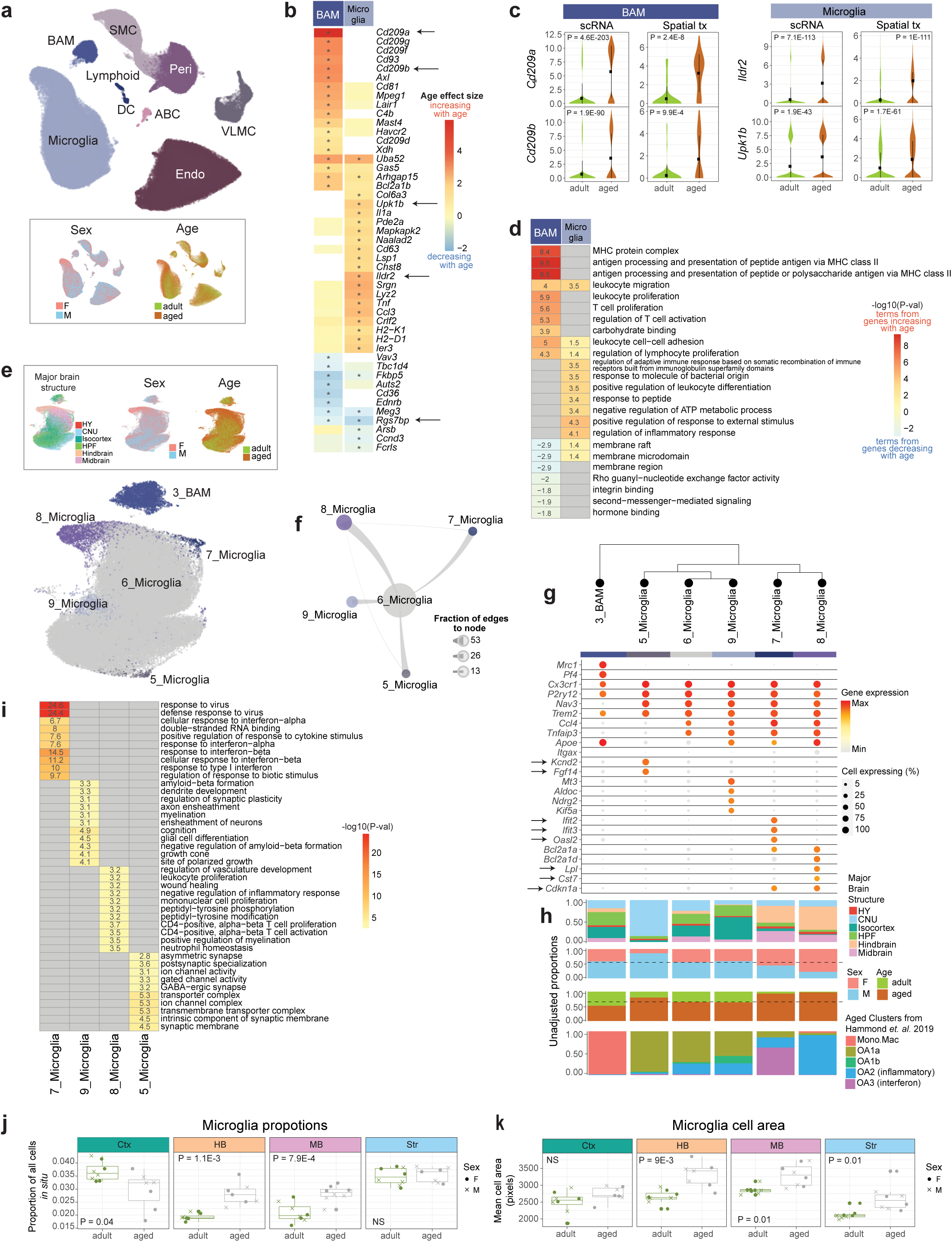
Age-associated changes in microglia and macrophages. **(a)** UMAP of all vascular and immune cell transcriptomes colored by subclass, sex, and age. **(b)** Heatmap of age effect sizes of top age-DE genes in BAM and microglia. Asterisk denotes statistical significance (see subclass level criteria in Methods). **(c)** Violin plot expression of *Cd209a* and *Cd209b* in BAM and *Ildr2* and *Upk1b* in microglia in scRNA-seq and spatial RSTE1 datasets. **(d)** Heatmap of the statistical significance of top GO terms enriched in top age-DE genes from BAM and microglia. Numbers in the plot represent -log_10_(p-value) of each term. Positive numbers are terms enriched in genes that increase with age and negative numbers are terms enriched in genes that decrease with age. **(e)** UMAP of immune cells including microglia and BAM, colored by cluster label, brain structure, sex, and age. **(f)** Constellation plot of microglia clusters colored by cluster created as described previously. **(g)** Marker gene expression in immune cell types organized in a dendrogram calculated from cluster DE genes. **(h)** Bar plot summaries for each cluster colored by brain structure, sex, age, and mapping label from Hammond *et al.* 2019 dataset. **(i)** Heatmap of statistical significance of top GO terms enriched in marker genes from non-homeostatic microglia clusters. **(j)** Changes in microglia created as in Figure 3e age calculated from spatial dataset RSTE1. **(k)** Changes in mean soma area of microglia cells with age as estimated from Baysor segmentation. Statistical significance for (j) and (k) are calculated with Student’s t-test. Each point represents a single replicate mouse sample.

In microglia with age, we observed upregulation of genes related to GO terms involving inflammatory response, response to bacteria, and others (**Figure 4d**). We also confirmed expression changes of genes observed by other single-cell studies of aging in microglia, including upregulation of *Ildr2* and *Upk1b* and downregulation of *Rgs7bp*^5,12,44^ with age (**Figure 4b,c**). *Upk1b* is a gene that encodes for uroplakin-1b and is included in the microglia “sensome”, a signature of genes expressed in microglia which encode proteins that sense endogenous ligands and microbes^45^. *Ildr2* is amongst GO terms related to protein localization to extracellular regions, which are enriched in genes that increase with age in microglia (**Supplementary Table 4**).

Upon further clustering of aged and adult brain immune cells, we identified 6 transcriptionally distinct clusters, 5 of which belong to microglia (**Figure 4e,f**). All microglia clusters expressed canonical microglia markers, including *Cx3cr1*, *P2ry12*, *Nav3*, and *Trem2* (**Figure 4g**). The largest microglia cluster (6_Microglia) contained 18,606 cells and was likely composed of the homeostatic microglia observed in both aged, adult, male, and female brains (**Figure 4h**). The four other microglia clusters were much smaller than cluster 6 (**Figure 4e,f**) and possibly represented different states of activated microglia. One of these clusters, cluster 5_Microglia, was very region and sex biased. It was found mostly in male CNU (specifically dorsal striatum) and uniquely expressed many genes including *Kcnd2* and proinflammatory *Fgf14* (**Figure 4g,h**). GO analysis revealed that genes involved in transporter and ion channel complex, as well as synapse related terms were amongst genes uniquely expressed in cluster 5_Microglia (**Figure 4i**).

We identified two age-biased clusters, 7_Microglia and 8_Microglia (**Figure 4h**). Both clusters show increased expression of the antiapoptotic Bcl-2 family members *Bcl2a1a*, and *Bcl2a1d*, which have been shown to increase in a variety of cell types with cell senescence^46^, as well as increased expression of cell senescence marker *Cdkn1a* (**Figure 4g**), consistent with prior studies detecting the accumulation of senescent microglia in aged mouse brain^47,48^. In addition, we found cluster-specific markers resembling those found by Hammond *et al.* in their scRNA-seq study profiling microglia throughout mouse lifespan^4^. Specifically, these authors found two age-enriched microglia clusters, OA2 and OA3, which expressed inflammatory markers and interferon-response genes, respectively^4^. By performing label transfer from their dataset to ours based on gene expression (**Methods**), we aligned our clusters 7_Microglia and 8_Microglia to Hammond’s OA3 and OA2 clusters, respectively (bottom bar of **Figure 4h**). We also found expression of similar cluster-specific genes in these two age-biased clusters, including increased expression of *Ifit2*, *Ifit3*, *Oasl2*, and other interferon-response genes in 7_Microglia, as well as increased expression of inflammatory markers such as *Cst7* and *Lpl* in cluster 8_Microglia, suggesting that these two clusters are likely the same cell types that were identified by Hammond *et al.* (**Figure 4g,h**). Of note, both these age-enriched clusters were mostly derived from hindbrain and midbrain. Marker genes for cluster 7 showed enrichment of GO terms related to interferon and virus response, while marker genes for cluster 8 showed enrichment of GO terms related to immune cell proliferation and activation (**Figure 4i**). Interferon signaling phenotypes were also observed in activated microglia from a mouse model of severe neurodegeneration^49^, suggesting the clusters we observe here may be precursors to microglia that are associated with neurodegenerative pathology.

Finally, to investigate whether proportions or size of microglia changed significantly with age throughout the brain, we estimated proportions and mean cell soma area (as estimated by segmentation) of microglia in 4 broad regions across the brain (**Figure 4j,k**) with spatial transcriptomics (dataset RSTE1). We found a significant increase in overall proportions of microglia in hindbrain and midbrain areas, no change in the striatum, and decrease in the cortex. We also observed an increase in the mean cell soma area of microglia in midbrain, hindbrain, and striatum, but not in the cortex (**Figure 4k**). These findings are partly consistent with prior findings of an increase in microglia counts with age in mouse VTA^50^, a decrease in microglia counts in mouse cortex^44^, and an increase in soma volume with age in microglia in the mouse somatosensory cortex^51^. However, overall, reports of changes in absolute numbers of microglia in rodents vary by region and study^44,51–53^. As such, our data support the idea that changes in microglia morphology and abundance with age vary by brain region.

### Changes in brain vascular cell types with age

Aging leads to loss of integrity and function of the brain microvasculature^54,55^. We characterized age-associated changes in the vascular cell subclasses found in our dataset, including arachnoid barrier cells (ABC; n = 546), vascular leptomeningeal cells (VLMC; n = 5,347), endothelial cells (n = 51,454), smooth muscle cells (SMC; n = 10,187), and pericytes (n = 17,187), which all display age-related DE genes (**Figure 2**). When plotted together in UMAP space, all vascular subclasses are transcriptionally highly distinct from one another (**Figure 4a**). Across these subclasses, endothelial cells showed the greatest number of age-DE genes, followed by pericytes, SMC, VLMC, and ABC (**Figure 2**). Due to the low number of ABCs in our dataset, we focus on the other 4 subclasses in the remainder of this section.

For endothelial cells, we found strong upregulation of *Hdac9* with age (**Extended Data Figure 7a**), and confirmed it by spatial transcriptomics (**Extended Data Figure 7b**). *Hdac9* gene and protein upregulation was previously observed in the ischemic brain and it exacerbates endothelial injury^56^, suggesting that normal endothelial cell function and thus oxygenation efficiency may be compromised in the brain with age. We also observed upregulation of many genes that encode proteins that are part of the MHC class I protein complex including *H2-Q7* and *H2-Q6*, as well as genes contributing to GO terms involving immune responses related to MHC class I upregulation and CD8 receptor binding (**Extended Data Figure 7a,c**). Together these findings suggest that there is an increase in antigen-presenting activity derived from intracellular proteins in endothelial cells with age. We also observed upregulation of similar MHC class I GO terms in VLMCs with age, although they appear to be driven by a different gene (*H2-D1*) (**Extended Data Figure 7a,c**).

VLMCs are fibroblast-like cells found in the brain. Across the VLMC subclass, we observed downregulation of genes that are involved in biomineralization and collagen extracellular matrix including collagens *Col11a1* and *Col3a1* (**Extended Data Figure 7a,c**), pointing to a decrease in structural integrity in this specialized cell type. Likewise, in SMC and pericytes, we observed downregulation of genes related to collagen extracellular matrix organization, although these changes were driven by different collagen genes, *Col4a1* and *Col4a2* (**Extended Data Figure 7a,c**). We confirmed downregulation of *Col4a2* in SMC and pericytes by spatial transcriptomics (**Extended Data Figure 7b**). Taken together, these results suggest loss of collagen expression and therefore, loss of extracellular matrix organization may be major contributors to the decreased structural integrity observed in brain vasculature with age. To assess potential changes in numbers of vascular cells with age, we calculated the proportion of each vascular cell type from spatial dataset RSTE1 (**Extended Data Figure 7d**). We found a significant decrease in the proportion of endothelial cells in the striatum, as well as a decrease in pericytes in the striatum and hindbrain regions. Interestingly, we observed an increase in the proportion of VLMCs in the hindbrain with age.

### Changes in astrocyte and ependymal cell class with age

Next, we investigated the Astro-Epen class of non-neuronal cells, which include telencephalic and non-telencephalic astrocytes (Astro-TE and Astro-NT, n = 143,167 and 118,221, respectively), astroependymal cells (n = 571), hypendymal cells (n = 164), tanycytes (n = 1,432), and ependymal cells (n = 2,923). When examining these cells in the UMAP space, we observed clear separation of the main Astro-TE and Astro-NT types by broad brain region, and the other smaller subclasses derived from specific brain regions as expected^17,57^ – for example, tanycytes were derived from the hypothalamus, whereas the ependymal cells came mostly from hindbrain and midbrain (**Figure 5a**). Across all subclasses found in the scRNA-seq dataset, tanycytes and ependymal cells showed the greatest numbers of age-DE genes (**Figure 2**). This was surprising, particularly given the relatively smaller cell numbers for these subclasses compared to the others (**Figure 5a**).

**Figure 5.**
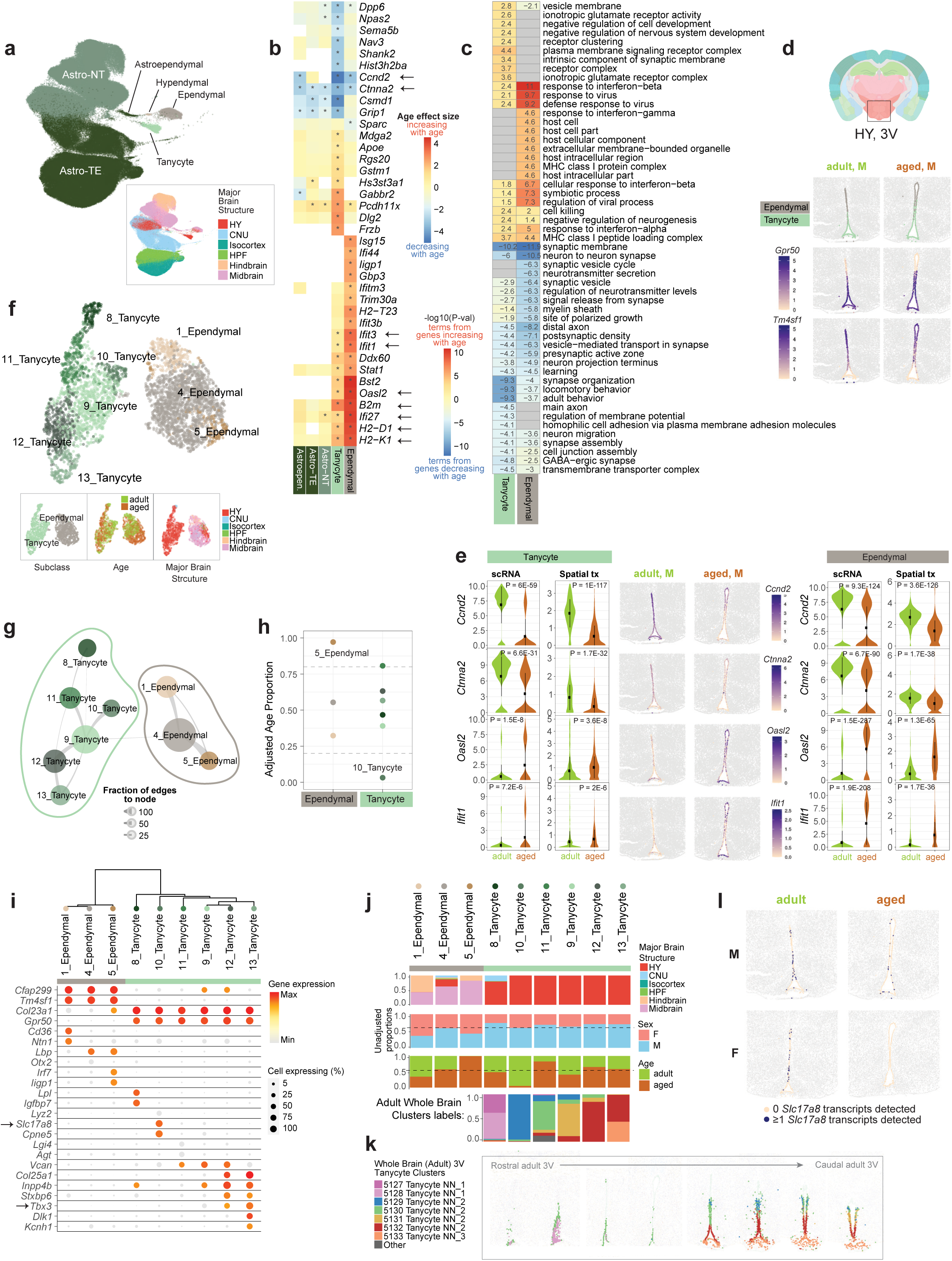
Age-associated changes in third ventricle tanycytes and ependymal cells. **(a)** UMAP of all Astro-Epen cell types colored by subclass and major brain structure. **(b)** Heatmap of age effect sizes of top age-DE genes in tanycytes and ependymal cells. Asterisk denotes statistical significance (see subclass level criteria in Methods). **(c)** Heatmap of the statistical significance of top GO terms enriched in top age-DE genes from tanycytes and ependymal cells. Numbers in the plot represent -log_10_(p-value) of each term. Positive numbers are terms enriched in genes that increase with age and negative numbers are terms enriched in genes that decrease with age. **(d)** Tanycyte and ependymal cell body locations in select samples from spatial dataset RSTE2, colored by subclass label (top), *Gpr50* (center), and *Tm4sf1* (bottom) expression. **(e)** Gene expression of *Ccnd2*, *Ctnna2*, *Oasl2,* and *Ifit1* across tanycytes (left) and ependymal cells (right) from scRNA-seq and spatial dataset RSTE2 represented by violin plots. Select adult and aged spatial RSTE2 samples are displayed in the center, colored by expression of each gene in tanycytes and ependymal cells. **(f)** UMAP of tanycytes and ependymal cell transcriptomes with additional adult cells from Yao *et al.* 2023 included, colored by cluster, subclass, age, and brain structure. **(g)** Constellation plot of clusters in (f), created as described previously. **(h)** Adjusted age proportion of each cluster from (g) colored by cluster and grouped by subclass. **(i)** Marker gene expression in tanycyte and ependymal cell clusters organized in a dendrogram calculated from cluster DE genes. **(j)** Bar plot summaries for each cluster colored by brain structure, sex, age, and adult cell label (see k) from Yao *et al.* 2023. **(k)** Location of tanycyte clusters in the Allen whole mouse brain cell type atlas^17^. **(l)** Visualization of *Slc17a8* gene expression changes in tanycytes and ependymal cells with age (*Slc17a8* gene expression was binarized in representative samples from spatial RSTE2 dataset).

Within the two main subclasses of astrocytes, Astro-TE and Astro-NT, we observed fewer age-DE genes (**Figure 2**). Furthermore, the types of age-DE genes differed between these two subclasses of astrocytes (**Extended Data Figure 8a,b**). In Astro-TE, there was an age-dependent downregulation of genes involved in neuron function-related terms such as axonogenesis and postsynaptic density, including *Dcc, Kcnd2,* and *Sema6d* (**Extended Data Figure 8b**; **Supplementary Table 4**). In Astro-NT, there was an age-dependent downregulation of genes involved in ion channel regulator activity, including *Kcnip4* and *Dpp6* (**Extended Data Figure 8b**; **Supplementary Table 4**). Using spatial transcriptomics, we found no significant change in astrocyte proportions with age, except for Astro-NT in the hindbrain region (**Extended Data Figure 8c**).

### Changes in third-ventricle tanycytes and ependymal cells with age

Ependymal cells are a type of ciliated glial cells that line the ventricles within the brain and the central canal of spinal cord. They assist in the circulation of cerebrospinal fluid throughout the ventricular system^58^. Tanycytes are a specialized form of ependymal cells that line the ventral and ventrolateral sides of the third ventricle (3V) in the hypothalamus and possess a single long protrusion that projects into the parenchyma of the hypothalamus^59^. Tanycytes are involved in regulating nutrient sensing and hormone signaling^59^. Tanycytes have also been shown to display adult neurogenic ability that may act as an adaptive mechanism in response to external factors such as physical activity and diet^60^. When we examined individual age-DE genes across these two subclasses, we found similar sets of age-DE genes and GO terms enriched with age across both subclasses, but not the other Astro-Epen subclasses (**Figure 5b, c**).

Using spatial transcriptomics, we clearly identified tanycytes and ependymal cells lining the third ventricle (dataset RSTE2, **Figure 5d**). We observed a dorsal-to-ventral transition between the two cell subclasses based on marker genes including *Gpr50* for tanycytes and *Tm4sf1* for ependymal cells (**Figure 5d**), allowing us to visually confirm and interrogate gene expression changes with age (center panels of **Figure 5e**).

Overall with age, there was an increase in many interferon response genes, such as *Ifi27*, *Ifit1, Ifit3,* and *Oasl2*, across ependymal cells, and to a fewer and less significant extent, in tanycytes (**Figure 5b; Supplementary Table 3**). There was also an increase in genes involved in the MHC class I response pathway, including *B2m*, *H2-K1* and *H2-D1*, across both ependymal cells and tanycytes (**Figure 5b; Supplementary Table 3**). These age-DE genes contributed to an enrichment of GO terms related to interferon-beta and virus responses, and MHC class I protein complex (**Figure 5c; Supplementary Table 4**). We confirmed increased expression of *Oasl2* and *Ifit1* with spatial transcriptomics (dataset RSTE2, **Figure 5e**).

Among the genes that decreased most strongly with age in both cell subclasses are the cell cycle gene *Ccnd2* and cadherin-associated protein gene *Ctnna2* (**Figure 5b,e**). *Ccnd2* has been shown to play an important role in adult neurogenesis^61^. *Ctnna2* is involved in the regulation of neuron migration and neuron projection development^62^. GO analysis revealed enrichment of terms related to neuronal structure and function in genes that were decreasing with age in both tanycytes and ependymal cells (**Figure 5c; Supplementary Table 4**). We also observed enrichment of terms related to negative regulation of neurogenesis and cell development in genes that were increasing with age (**Figure 5c; Supplementary Table 4**), which may suggest a decrease in neurogenic potential in tanycytes with age.

To investigate changes with age at the finer cell-type level, we further clustered both tanycytes and ependymal cells. Because our original tanycyte scRNA-seq dataset was unbalanced towards a larger number of aged cells, we included additional cells from the adult whole mouse brain dataset^17^ that were originally excluded because they came from a slightly different dissection region (**Methods**). After clustering, we defined 6 tanycyte and 3 ependymal clusters (**Figure 5f,g**). Three ependymal clusters displayed unique gene markers (**Figure 5i**) and came from different regions of the brain, with cluster 1_Ependymal found in both midbrain and hindbrain, 4_Ependymal found in mostly midbrain and hypothalamus, and 5_Ependymal mostly found in midbrain (**Figure 5f,j**). After calculating the adjusted age proportion, we found that one of these ependymal clusters (5_Ependymal) consisted almost entirely of aged cells, and as such, we consider this cluster age-biased (**Figure 5h,j**). Unique marker genes for this cluster include interferon response genes *Iigp1* and *Irf7* (**Figure 5i**), further supporting increased interferon signaling with age in ependymal cells.

The six tanycyte clusters all displayed unique sets of marker genes (**Figure 5i**) mostly aligning with different known types of tanycytes^59,63^. To estimate the spatial location of each tanycyte cluster, we examined cluster labels from the thoroughly annotated adult tanycyte cells and their location on the corresponding Allen whole mouse brain spatial atlas^17^ (**Figure 5j,k**). We found representation of nearly all adult whole brain tanycyte clusters: 8_Tanycyte represents tanycytes from rostral 3V, 10_Tanycyte represents the most dorsal α1 subtype (aligned with the dorsomedial and ventromedial nuclei of the hypothalamus, DMH and VMH), 9_Tanycyte and 11_Tanycyte represent α2 subtypes (aligned with dorsal ARH) which are ventral to α1, and 12_Tanycyte and 13_Tanycyte represent the most ventral tanycyte subtypes, β1 (aligned with ventral ARH) and β2 (aligned with the median eminence, ME), respectively (**Figure 5j,k**).

Amongst the tanycyte clusters, we observed one cluster that appeared to be adult-biased, cluster 10_Tanycyte (**Figure 5h)**, likely the cluster representing α1 tanycytes (**Figure 5j,k**). Marker genes for cluster 10_Tanycyte include *Slc17a8* and *Cpne5* (**Figure 5i**). We also confirmed decreased expression of *Slc17a8* in the dorsal tanycytes of the 3V in the spatial data (**Figure 5l**). *Slc17a8* is regarded as a marker for α1 tanycytes^63^, so loss of *Slc17a8* with age suggests that tanycyte types may become less distinctive with age.

### Changes in hypothalamic *Tbx3*+ neurons with age

Across the neuronal subclasses identified in our dataset, those with the greatest numbers of age-DE genes were hypothalamic neurons (**Figure 2**). There were four classes of hypothalamic neurons in our dataset, including HY GABA, HY Glut, CNU-HYa Glut, and HY MM Glut (MM standing for medial mammillary nucleus), which were confirmed by *Slc32a1* and *Slc17a6* expression (**Figure 6a**). Under these classes, there were 29 subclasses that displayed unique marker gene expression (**Extended Data Figure 2**, **Figure 6b, Supplementary Table 2**; neuronal subclass names were transferred from the Allen Mouse Whole Brain Atlas^17^, where they were named for the most dominant brain region localization and transcription factor expression), altogether capturing the vast cell type complexity we previously reported in the adult mouse hypothalamus^17^.

**Figure 6.**
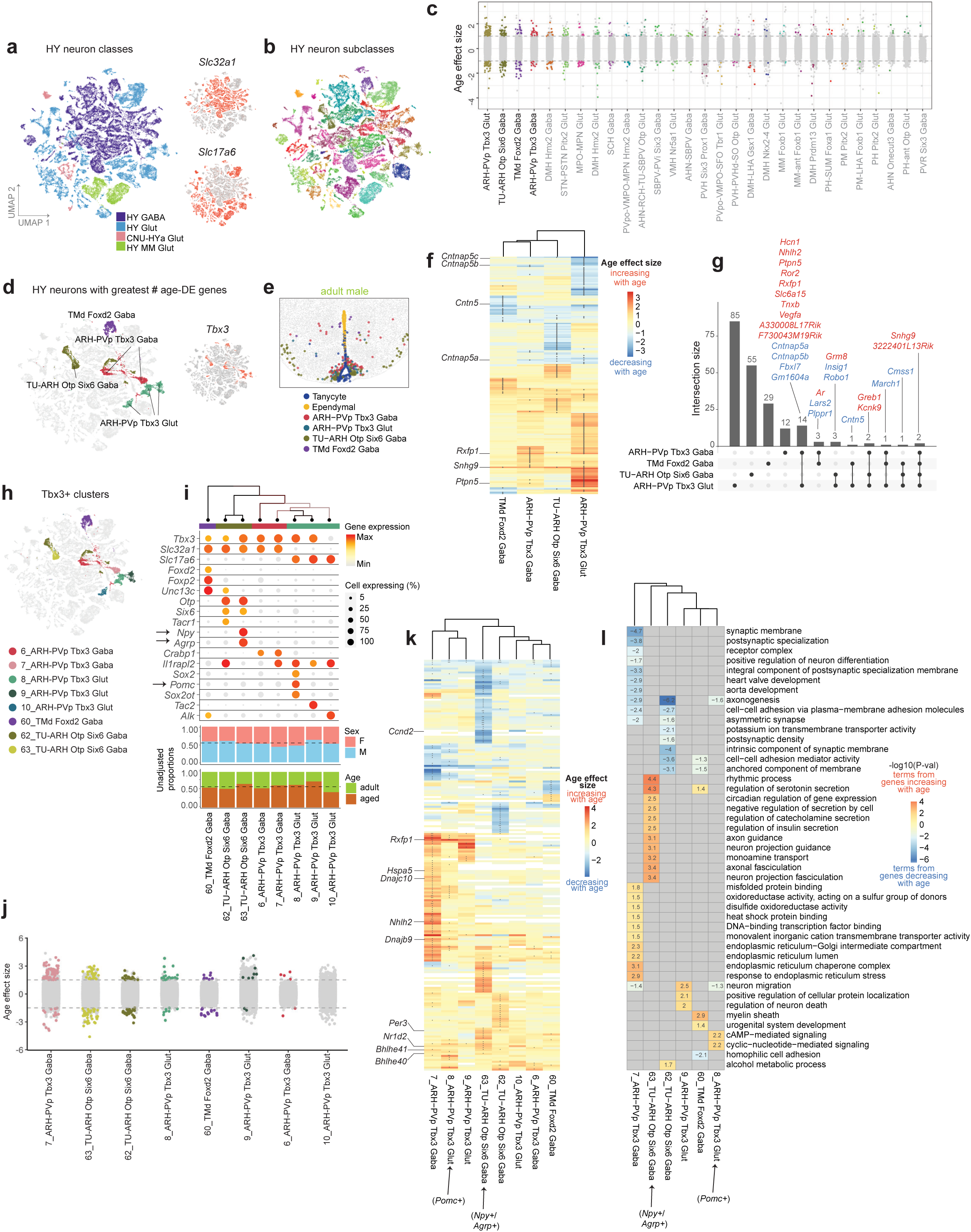
Age-associated changes in *Tbx3*+ hypothalamic neurons. **(a-b)** UMAP of all hypothalamic (HY) neurons colored by (a) class, *Slc32a1* and *Slc17a6* expression, and (b) subclass. **(c)** Age effect sizes of age-DE genes from hypothalamic neuronal subclasses ordered by the number of age-DE genes, with significant age-DE genes colored. Labels for the top 4 subclasses are emphasized with darker font on the left. **(d)** Subclasses with the greatest numbers of age-DE genes highlighted and *Tbx3* expression shown in the same UMAP space as (a). **(e)** Neurons, tanycyte and ependymal cell body locations in a representative sample from spatial dataset RSTE2 demonstrating colocalization of subclasses from (d) around the third ventricle. **(f)** Heatmap of age effect sizes of all age-DE genes in *Tbx3*+ neuronal subclasses. Asterisks denote statistical significance. Dendrogram represents hierarchical clustering of subclasses based on age effect sizes. Genes discussed in text are labeled. **(g)** Upset plot of overlapping age-DE genes between the four *Tbx3*+ neuronal subclasses. Genes colored in red increase with age while genes colored in blue decrease with age in scRNA-seq data. **(h)** *Tbx3*+ neuronal clusters colored in the same UMAP space as (a). **(i)** Marker gene expression in *Tbx3*+ neuronal clusters organized in a dendrogram calculated from cluster DE genes. Bar plot summaries of each cluster colored by sex and age are below. **(j)** Age effect sizes of age-DE genes from *Tbx3*+ clusters ordered from the greatest to least number of age-DE genes, with significant age-DE genes colored. **(k)** Heatmap of age effect sizes from all age-DE genes from *Tbx3*+ clusters. Asterisks denote statistical significance (Methods). Dendrogram represents hierarchical clustering of clusters based on age effect sizes. Genes discussed in text are labeled. **(l)** Heatmap of statistical significance of top GO terms enriched in marker genes from all *Tbx3*+ neuronal clusters.

Across the 29 hypothalamic neuronal subclasses, the subclasses with the greatest numbers of age-DE genes were ones associated with hypothalamic regions proximal to the third ventricle, including the arcuate nucleus (ARH), posterior periventricular nucleus (PVp), dorsal tuberomammillary nucleus (TMd), and dorsomedial nucleus (DMH) (**Figure 6c**). Remarkably, the 4 subclasses with the greatest numbers of age-DE genes, i.e., ARH-PVp Tbx3 Glut (n = 1,134 cells), TU-ARH Otp Six6 Gaba (n = 1,191), TMd Foxd2 Gaba (n = 711), and ARH-PVp Tbx3 Gaba (n = 1,031), all had highly specific expression of the transcription factor *Tbx3* (**Figure 6d**). Interestingly, we also observed distinctive *Tbx3* expression in ventral tanycytes, but not in the more rostrally and dorsally located tanycytes (**Figure 5i**).

The cell bodies of these four subclasses were all located directly proximal to the third ventricle, with the ARH subclasses interacting directly with the ventral β-type tanycytes (spatial dataset RSTE2; **Figure 6e**). These four *Tbx3* positive (*Tbx3*+) subclasses also demonstrated highly distinct signatures of aging, as reflected by the different sets of age-DE genes (**Figure 6f**) that contained subsets of age-DE genes either unique to each subclass or shared among multiple or all subclasses (**Figure 6g**). All four subclasses demonstrated an increase in *Snhg9*, a non-coding small nucleolar RNA host gene that has bene implicated in the development of obesity^64^ and as a biomarker for various cancers^65,66^. We observed downregulation of many genes coding for cell-adhesion contactin and contactin associated proteins, specifically of family member 5 (*Cntn5, Cntnap5a, Cntnap5b, Cntnap5c*), across one or more subclasses. We also observed an increase in *Ptpn5* with age, a biomarker of many neurodegenerative and neuropsychiatric disorders including Alzheimer’s, Parkinson’s, Huntington’s, schizophrenia, and others^67^.

Next, we investigated these *Tbx3*+ neurons at the cluster level. Using *de novo* clustering, we split these four subclasses into the following sets of clusters (**Figure 6h**): 3 ARH-PVp Tbx3 Glut clusters (labeled as clusters 8, 9, and 10), 2 ARH-PVp Tbx3 GABA clusters (clusters 6 and 7), and 2 TU-ARH Otp Six6 Gaba clusters (clusters 62 and 63). TMd Foxd2 Gaba cells remained as one population and were not split into additional clusters. Each cluster was relatively balanced in age and sex distributions and displayed unique expression of combinations of marker genes, including expression of namesake transcription factors *Tbx3*, *Otp*, *Six6*, and *Foxd2* (**Figure 6i**). Different clusters within each subclass exhibited unique sets of DE genes related to age. Additionally, specific clusters within a subclass appeared to predominantly contribute to the age-associated gene expression changes observed at the subclass level (**Figure 6j, k**). For example, between the two ARH-PVp Tbx3 Gaba clusters, cluster 7 demonstrated the greatest number of age-DE genes across all *Tbx3*+ clusters, while cluster 6 had far fewer age-DE genes. Similarly, among the 3 ARH-PVp Tbx3 Glut clusters, most age-associated changes were observed in clusters 8 and 9, but not 10. Interestingly, hierarchical clustering based on age effect sizes of the top age-DE genes across clusters grouped clusters 7, 8, and 9 in one branch, suggesting that despite being from different Glut and GABA subclasses, these 3 clusters appear to age more similarly than other *Tbx3*+ clusters (**Figure 6k**).

Neurons in the ARH are known for, among many functions, the critical role they play in modulation of energy homeostasis. For example, the well-characterized agouti-related peptide (AgRP) and proopiomelanocortin (POMC) neurons stimulate or inhibit food intake, respectively^68,69^ and are among the neuronal types that show the greatest numbers of gene expression changes under diet perturbation, including fasting and high fat diets^70^. AgRP neurons are characterized by expression of *Npy* and *Agrp*, while POMC neurons are characterized by expression of *Pomc*. In our *Tbx3*+ clusters, cluster 63_TU-ARH Otp Six6 Gaba shows highly specific expression of *Npy* and *Agrp*, while cluster 8_ARH-PVp Tbx3 Glut shows specific expression of *Pomc* (**Figure 6i**), suggesting these two clusters may participate in the canonical neuronal circuit that regulates food intake.

When we performed GO analysis on cluster age-DE genes, we found enrichment of genes related to cAMP-mediated signaling in *Pomc*+ cluster 8, a pathway implicated in many biological processes, including anti-aging pathways^71,72^ (**Figure 6l; Supplementary Table 4**). We also observed significant increase in expression of *Rxfp1* with age (**Figure 6k; Supplementary Table 3**), a gene encoding a G-protein coupled receptor that binds the highly evolutionarily conserved peptide relaxin-3 that mainly signals through the cAMP pathway^73^. Relaxin-3, which is encoded by the gene *Rln3,* is involved in various physiological processes such as feeding, arousal, stress response, and cognition. It is widely distributed throughout the brain as well as peripheral tissues^74^. We also observed increased expression of *Rxfp1* with age in cluster 7_ARH-PVp Tbx3 Gaba, as well as at the subclass level in both ARH-PVp Tbx Glut and GABA types, suggesting that clusters 7 and 8 are driving the increase in *Rxfp1* at the subclass level. In cluster 7, we observed significant enrichment of upregulated endoplasmic reticulum-localized heat shock protein genes, including *Hspa5, Dnajb9,* and *Dnajc10* (**Figure 6k,l; Supplementary Table 4**), an aging signature that appears to be specific to this cluster only. Furthermore, in cluster 7, the age-DE gene with the strongest age effect size was *Nhlh2*, which was also uniquely changing with age only in cluster 7 (**Figure 6k**). *Nhlh2* is a transcription factor that has been implicated in regulating processes related to obesity and fertility^75^.Amongst genes increasing with age in the *Agrp*+ cluster 63_ TU−ARH Otp Six6 Gaba, we found enrichment of terms related to monoaminergic neurotransmitter secretion and circadian regulation of gene expression (**Figure 6l; Supplementary Table 4**). Included in the circadian and rhythmic process related genes, we observed *Bhlhe40, Bhlhe41, Nr1d2,* and *Per3* increasing with age only in the *Agrp*+ cluster (**Figure 6k; Supplementary Table 3**), suggesting that temporal and rhythmic control of behaviors like feeding, a known function of *Agrp*+ neurons^76^, may become altered with age. Amongst genes uniquely decreasing with age in cluster 63 was *Ccnd2*, which we also observed decreasing in tanycytes and ependymal cells (**Figure 5b; Extended Data Figure 5b**). Taken together, we find that there are strikingly diverse differences in cluster-level aging signatures in *Tbx3*+ hypothalamic neurons, even within the same subclass, lending additional credence to a single-cell approach for investigating age-specific changes across cell types in the brain.

## Discussion

A gradual loss of homeostasis across many aspects of cellular and organismal function occurs with aging. Many of these themes, or hallmarks, of aging, including genomic instability, epigenetic alteration, chronic inflammation, cellular senescence, deregulated nutrient-signaling, etc., have been observed in multiple invertebrate and vertebrate species^2,3^. However, the mechanisms that govern systemic aging at the organismal level across complex tissue types and organ systems remain unclear. Certain cell types are more vulnerable to specific aspects of aging than others, and likely communicate and interact with other cell and tissue types to integrate both intrinsic and extrinsic signals that ultimately contribute to decline in cellular and organismal health. As such, a single-cell approach to characterizing transcriptional changes in the brain-wide neural network is a critical step towards fully understanding brain-wide, and eventually, organismal aging.

In this study, we present a large-scale, comprehensive single-cell transcriptomic atlas and comparative analysis of the young adult and aged mouse brains. Large cell numbers, high quality of transcriptomes, brain-wide coverage, and detailed annotation of cell types using our newly created Allen whole mouse brain cell types atlas^17^ enabled us to precisely pinpoint the regions and cell types in the brain that may be particularly vulnerable to aging. We find evidence for conservation of many of the canonical hallmarks of aging across various cell types within the aged mouse brain. This includes 1) increased expression of cell senescence markers in age-enriched oligodendrocyte and microglia clusters (**Figure 3, 4**), 2) increased systemic inflammation as suggested by the identification of age-enriched proinflammatory microglia clusters, 3) oligodendrocyte clusters with increased inflammation signals and recruitment of microglia, 4) ependymal clusters with increased interferon signaling (**Figure 3-5**), 5) decrease in new myelination as indicated by the depletion of immature oligodendrocyte cell types in the aged brain (**Figure 3**), and 6) decrease of structural integrity in the brain vasculature as indicated by the downregulation of extracellular matrix genes in the smooth muscle and endothelial cell types (**Extended Data Figure 6**). Interestingly, many of these changes are found to be more pronounced in hindbrain and midbrain regions. Although not investigated in detail here, we also observe signs of deterioration of neuronal function with aging, including altered gene expression in a number of cortical and hippocampal neuronal types (**Figure 2**), changes in immature neuronal types that are involved in adult neurogenesis (**Figure 2**), as well as potentially altered neuron-astrocyte interactions (**Extended Data Figure 8**). Most prominently, we observe evidence of altered regulation of nutrient-sensing and energy homeostasis via many gene expression changes in tanycytes, ependymal cells, and *Tbx3*+ neurons localized around the arcuate nucleus and third ventricle of the hypothalamus, site of the canonical melanocortin circuit of the brain that regulates energy homeostasis (**Figure 5, 6**).

Deregulated nutrient sensing and the gradual loss of energy homeostasis is one of the most extensively investigated aspects in aging and longevity research. Moreover, caloric restriction and intermittent fasting have been shown to delay aging-associated structural and functional decline and increase longevity across several animal species^77^. The somatotrophic axis – one of the most highly conserved signaling axis observed over evolution – involves growth hormone (GH)-mediated stimulation of insulin growth factor and mammalian target of rapamycin (MTOR) signaling network, manipulation of which increases lifespan and health span across all organisms tested^78,79^.

The area surrounding the third ventricle of the hypothalamus, including the arcuate nucleus, is commonly regarded as one of the circumventricular organs of the brain: it contains a more permissive blood vascular system than the rest of the brain, allowing nutrients and hormones from blood to interact more freely with neurons and glia in that region^80^. MTOR activity increases during aging in hypothalamic neurons, contributing to age-related obesity, which is reversed by direct infusion of rapamycin to the hypothalamus^81^. In addition to the MTOR pathway, the ALK signaling pathway, another nutrient-sensing pathway, is induced in the hypothalamus by feeding^82^, and hypothalamus-specific deletion of *Alk* in mice promotes resistance against diet-induced obesity, a common age-associated phenotype^82^.

We find that *Tbx3*+ cell types in the hypothalamus, both neurons and tanycytes, may be more susceptible to age-related changes than other cells in the brain. We observe highly diverse gene expression changes among these cell types that are concentrated around the 3^rd^ ventricle (**Figure 6**), suggesting differential roles these cell types play and their complex interactions in the aging process. As of yet, we do not know whether these changes are driven by cellular programs that are protective against or susceptible to aging, or both. There is evidence to suggest that in mouse embryonic fibroblasts, *Tbx3* expression may suppress cell senescence^83^, a key contributor to cellular aging. *Tbx3* is also differentially expressed at high levels in many enteric neurons that govern the function of the gastrointestinal tract^84^, suggesting that there may be common expression patterns between hypothalamus and the enteric nervous system that may be relevant to metabolic homeostasis and aging. In addition to many hypothalamic neurons, tanycytes are also regarded as a key integrator of nutrient and sex hormone signaling within the brain^59^. Tanycytes have also demonstrated adult neurogenic and gliogenic ability, possibly in response to changes in diet^85^.

Given the proximity of both tanycytes, ependymal cells, and Tbx3+ neurons to the third ventricle, our results suggest that cells surrounding the third ventricle in the hypothalamus, may represent a critical focal point of the accumulation of age-associated changes in the brain. Furthermore, the highly conserved role POMC and AgRP neurons play in appetite regulation and energy homeostasis, as well as the role tanycytes play in nutrient sensing, coupled with the extensive body of literature implicating nutrient dysregulation in aging biology^86^ suggest that this region of the brain may act as a key systemic integrator of nutrient and energy signaling across the entire organism that heavily influences cellular and/or organismal aging.

The dataset we present here represents the most extensive and comprehensive transcriptomic analysis of the normal aged mouse brain that we know of to date. The identification of a variety of robust and highly significant gene expression changes with aging across many neuronal and non-neuronal cell types throughout the brain demonstrates the power and necessity of single-cell approaches to revealing the mechanisms that govern complex systemic phenotypes like aging. The results and insights from this work will serve as a foundational resource for the neuroscience and aging research communities to facilitate detailed investigation of age-associated phenotypes in the brain and the body and the interaction between aging and various diseases.

## Methods

### Mouse breeding and husbandry

All procedures were carried out in accordance with Institutional Animal Care and Use Committee protocols at the Allen Institute for Brain Science. Mice were provided food and water *ad libitum* and were maintained on a regular 14:10 hour day/night cycle at no more than five adult animals of the same sex per cage. Mice were maintained on the C57BL/6J background. We excluded any mice with dermatitis, anophthalmia, microphthalmia, seizures, or abdominal masses.

We used 44 aged mice (20 female, 22 male) and 52 adult mice (25 female, 27 male) to collect 2,777,165 cells for 10xv3 scRNA-seq. All adult animals were also included in the Allen whole mouse brain cell type atlas^17^. Aged animals were euthanized at P540-553 (approximately 18 months) and adult animals were euthanized at P53-69 (approximately 2 months). No statistical methods were used to predetermine sample size. All donor animals used in this study are listed in **Supplementary Table 1.**

We isolated a total of 272 libraries from 96 animals – each animal contributed 1-6 libraries. All libraries are listed in **Supplementary Table 1**. Transgenic driver lines were used for fluorescence-positive cell isolation by FACS to enrich for neurons. Approximately half the libraries (n = 133) were sorted for neurons from the pan-neuronal *Snap25-IRES2-Cre* line (JAX strain #023525) crossed to the *Ai14*-tdTomato reporter (JAX strain #007914) ^87,88^ (**Supplementary Table 1**). For unbiased sampling without FACS, we used either *Snap25-IRES2-Cre/wt;Ai14/wt* mice*, Ai14/wt* mice, or in very few cases wildtype C57BL/6J mice. The transgenic *Snap25-IRES2-Cre* line was backcrossed to C57BL/6J for at least 10 generations before crossing and can be considered congenic. The transgenic *Ai14* line was backcrossed to C57BL/6J for at least 5 generations before crossing and can be considered incipient congenic.

### 10X single-cell RNA sequencing

#### Single-cell isolation

We used the Allen Mouse Brain Common Coordinate Framework version 3 (CCFv3; RRID: SCR_002978) ontology^21^ (http://atlas.brain-map.org/) to define brain regions for profiling and boundaries for dissection. We covered all regions of the brain by sampling at top-ontology level with judicious joining of neighboring regions. These choices were guided by the fact that microdissections of small regions are difficult. Therefore, joint dissection of neighboring regions was sometimes necessary to obtain sufficient numbers of cells for profiling.

Single cells were isolated by adapting previously described procedures^16,89^. The brain was dissected, submerged in ACSF, embedded in 2% agarose, and sliced into 350-μm coronal sections on a compresstome (Precisionary Instruments). Block-face images were captured during slicing. Regions of interest (ROIs) were then microdissected from the slices and dissociated into single cells as previously described^16,89^. Fluorescent images of each slice before and after ROI dissection were taken at the dissection microscope. These images were used to document the precise location of the ROIs using annotated coronal plates of CCFv3 as reference.

Dissected tissue pieces were digested with 30 U/ml papain (Worthington PAP2) in ACSF for 30 minutes at 30°C. Due to the short incubation period in a dry oven, we set the oven temperature to 35°C to compensate for the indirect heat exchange, with a target solution temperature of 30°C. Enzymatic digestion was quenched by exchanging the papain solution three times with quenching buffer (ACSF with 1% FBS and 0.2% BSA). Samples were incubated on ice for 5 minutes before trituration. The tissue pieces in the quenching buffer were triturated through a fire-polished pipette with 600-µm diameter opening approximately 20 times. The tissue pieces were allowed to settle and the supernatant, which now contained suspended single cells, was transferred to a new tube. Fresh quenching buffer was added to the settled tissue pieces, and trituration and supernatant transfer were repeated using 300-µm and 150-µm fire polished pipettes. The single cell suspension was passed through a 70-µm filter into a 15-ml conical tube with 500 µl of high BSA buffer (ACSF with 1% FBS and 1% BSA) at the bottom to help cushion the cells during centrifugation at 100 x g in a swinging bucket centrifuge for 10 minutes. The supernatant was discarded, and the cell pellet was resuspended in the quenching buffer. We collected 1,508,284 cells without performing FACS. The concentration of the resuspended cells was quantified, and cells were immediately loaded onto the 10x Genomics Chromium controller.

To enrich for neurons or live cells, cells were collected by fluorescence-activated cell sorting (FACS, BD Aria II) using a 130-μm nozzle. Cells were prepared for sorting by passing the suspension through a 70-µm filter and adding Hoechst or DAPI (to a final concentration of 2 ng/ml). Sorting strategy was as previously described^16,17^, with most cells collected using the tdTomato-positive label. 30,000 cells were sorted within 10 minutes into a tube containing 500 µl of quenching buffer. We found that sorting more cells into one tube diluted the ACSF in the collection buffer, causing cell death. We also observed decreased cell viability for longer sorts. Each aliquot of sorted 30,000 cells was gently layered on top of 200 µl of high BSA buffer and immediately centrifuged at 230 x g for 10 minutes in a centrifuge with a swinging bucket rotor (the high BSA buffer at the bottom of the tube slows down the cells as they reach the bottom, minimizing cell death). No pellet could be seen with this small number of cells, so we removed the supernatant and left behind 35 µl of buffer, in which we resuspended the cells. Immediate centrifugation and resuspension allowed the cells to be temporarily stored in a high BSA buffer with minimal ACSF dilution. The resuspended cells were stored at 4°C until all samples were collected, usually within 30 minutes. Samples from the same ROI were pooled, cell concentration quantified, and immediately loaded onto the 10x Genomics Chromium controller.

#### cDNA amplification and library construction

For 10x v3 processing, we used the Chromium Single Cell 3′ Reagent Kit v3 (1000075, 10x Genomics). We followed the manufacturer’s instructions for cell capture, barcoding, reverse transcription, cDNA amplification and library construction. We targeted a sequencing depth of 120,000 reads per cell; the actual average achieved was 80,118 ± 35,612 (mean ± SD) reads per cell across 272 libraries (**Supplementary Table 1**).

#### Sequencing data pre-processing

All libraries were 10xv3 samples and processed as previously described^16,17^. All libraries were sequenced on Illumina NovaSeq6000 and sequencing reads were aligned to the mouse reference (mm10/gencode.vM23) using the 10x Genomics CellRanger pipeline (version 6.0.0) with the *–include introns* argument to include intronicaly mapped reads.

To remove low quality cells, we used a stringent QC process. Cells were first filtered by a broad set of quality cutoffs based on gene detection, qc score, and doublet score. As we previously described^17^, the qc score was calculated by summing the log-transformed expression of a set of genes, whose expression level is decreased significantly in poor quality cells. Briefly, these are housekeeping genes that are strongly expressed in nearly all cells with a very tight co-expression pattern that is anti-correlated with the nucleus-enriched transcript *Malat1*. We use this qc score to quantify the integrity of cytoplasmic mRNA content. Doublets were identified using a modified version of the **DoubletFinder** algorithm^90^. For this preliminary round of filtering, we included cells with gene detection > 1000, qc score > 50, and doublet score < 0.3. Using these thresholds, 1,999,976 cells remained in the dataset (**Extended Data Fig 1a**).

#### Clustering single cell RNA-seq data

Following the initial round of filtering described above, adult and aged single-cell transcriptomes were co-clustered over two rounds of clustering. The goal for the first round of clustering was to assign a cell class identity to every unlabeled (aged) cell and filter out low-quality (noise) clusters. The goal of the second round of clustering was to assign a subclass identity to every unlabeled (aged) cell and filter out additional low-quality clusters. All adult cells in the dataset already had labels because they are also part of the Allen whole mouse brain cell type taxonomy^17^. For both rounds, clustering was performed independently with the in-house developed R package **scrattch.bigcat** as was previously described^17^ (available via github https://github.com/AllenInstitute/scrattch.bigcat),. This package is version of R package **scrattch.hicat**^16^ that can cluster large datasets. Detailed functionality of scrattch.bigcat was discussed in our previous paper^17^. We used the automatic iterative clustering method, *iter_clust_big*, to peform clustering in a top-down manner into cell types of increasingly finer resolution. This method performs clustering without human intervention, while ensuring that all pairs of clusters, even at the finest level, were separable by differential gene expression criteria (*q1.th = 0.4, q.diff.th = 0.7, de.score.th = 300, min.cells = 50)* for both rounds of clustering. Following each round of clustering using *iter_clust_big*, we used the function *merge_cl* to merge clusters based on total number and significance of shared DE genes. For round 1, the criteria used for merge_cl were identical to those previously described for clustering. For round 2, the criteria used for merge_cl were almost identical with the exception of increasing *min.cells = 100*.

#### Assigning labels to aged cells and removing low-quality clusters

We observed 2,467 clusters after the first round of clustering. At this point, all cells were assigned a cell category (Glut, GABA, Dopa, Sero, IMN or NN). Since the adult cells have been previously published and annotated^17^, cell annotations for aged cells were assigned based on cluster membership with annotated adult cells. Specifically, clusters that contained >5% of annotated adult cells were assigned that cell category. Category-labeled clusters were then filtered based on cell category-specific cluster-level thresholds (**Supplementary Table 5**, **Extended Data Fig 1a**). Clusters with >80% contribution from a single library were also filtered out to minimize donor bias in the final dataset. Clusters with <5% adult cells were retained in the dataset and carried over into the next round of clustering. Since adult cells that were previously deemed to be low quality^17^ were also included in clustering, clusters with the majority of low-quality cells were also filtered out. In total, 1,197 clusters were removed based on these criteria after the first round of clustering (n = 779,838 cells removed). This resulted in the dataset of 1,220,138 cells, which were carried over into the second round of clustering (**Extended Data Fig 1a**).

After the second round of clustering, we observed 928 clusters. All clusters were then assigned subclass identities in a process similar to that described above. Clusters with <5% adult cells were now mapped directly to the Allen whole mouse brain cell type taxonomy^17^ (see “Label transfer via mapping” section below) and entire clusters were assigned to the most common subclass within the group of cells that made up that cluster. Annotated clusters were then filtered using class-level quality metrics and other quality metrics similar to those in the above paragraph (**Supplementary Table 5**, **Extended Data Fig 1a**). After this second round of cluster-level filtering, 31 clusters were removed (n = 34,934 cells removed) and 1,185,204 cells remained in the dataset. Remaining cells and resultant subclass annotations were used for all downstream analysis (**Extended Data Fig 1a**).

#### Label transfer via mapping

For assigning identities of cells in clusters with >95% aged cells, we mapped them to a reference taxonomy as previously described^17^. Briefly, we assigned their cell type identities by mapping them to the nearest cluster centroid in the reference taxonomy using the corresponding Annoy index as implemented in the R package **scrattch.mapping**. We also used this approach for assigning cell type identities for cells segmented from Resolve spatial data to the Allen whole mouse brain cell type taxonomy^17^ or external datasets as reference, using different gene lists based on the contexts. For mapping to the oligodendrocyte dataset from Marques *et. al.*^30^, we used a list of 195 genes. For mapping to the microglia dataset from Hammond *et. al.*^4^, we used a list of 72 genes. For both external datasets, gene lists were assembled based on prominent marker genes from each external reference cluster. When mapping confidence score was needed, we sampled 80% genes from the marker list randomly, and performed mapping 100 times. We define the fraction of times a cell is assigned to a given cell type as the mapping probability to that type.

#### Identifying age-associated DE genes

Age-associated DE genes were calculated using the R package **MAST**^22^, a widely used statistical framework designed for modeling biological effects from scRNA-seq data. Briefly, MAST fits a two-part generalized linear model and also allows for adaptive thresholding of gene expression data to account for dropout rate. Upon inspection using MAST’s thresholdSCRNACountMatrix function, we found that for most cases, genes expressed at a frequency of at least 10% did not reveal many genes with non-zero bimodal bins, so we did not implement any adaptive thresholding in our DE gene analysis.

DE genes were calculated at the subclass, supertype, and cluster level. For all tests, only genes that were expressed at a frequency of >10% were tested (i.e., only genes expressed in at least 10% of query cells were included). Only subclasses with at least 50 aged and 50 adult cells were evaluated for DE genes. To decrease running time, for large subclasses, we subsampled them to a maximum of 1,000 cells per age.

At the subclass level, we used the following two statistical models to model they effect of age on gene y including various covariates:

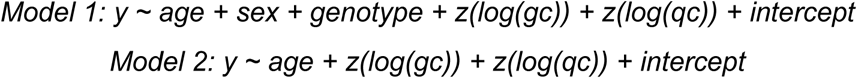

where age, sex, and genotype are all categorical variable with 2, 2, and 3 levels, respectively, and gene detection (gc) and QC score (qc) are log transformed and then z-score normalized. We included both gene detection and QC score in each model to account for potential effects that various FACS population plans had on library quality (**Extended Data Figure 4a**). A likelihood ratio test was computed between each model with and without the age term to generate p-values. These p-values were corrected for multiple hypothesis testing with the Bonferonni correction. The effect size estimate for the age term for each model can be interpreted as the log_2_-fold change (logFC) of each gene. However, due to the additional covariates, logFC estimated by the models often varied widely from those calculated without covariate adjustment. As such, we refer to this term as “age effect size” throughout the main body of the text, rather than logFC.

Since age effect sizes estimated by these two models differed widely for certain cell types, particularly smaller neuronal populations, we chose to consider a gene significant if and only if it exceeded statistical cutoffs (p < 0.01 & age effect size > 1 or < −1) for both Model 1 and 2. For all figures that plot heatmaps of age effect sizes of subclass age-DE genes, age effect sizes from Model 1 were used. At the supertype and cluster level, only results from Model 1 are presented.

For the vast majority of age-DE genes presented here, the directionality of age effect sizes between the two models agrees with one another. However, for a very small number of genes (6 out of 1,253 unique genes), the directionality disagrees, with most of these being changes in expression of the X-inactivation gene *Xist* across various hypothalamic neuron types (**Extended Data Figure 5b; Supplementary Table 3**) which may be due to the imbalance between libraries of different FACS population plans, sex, and age (**Extended Data Figure 4a**). However, as a recent study showed that *Xist* expression increases in aged female hypothalamic neurons^11^, in all figures, we display the age effect size of the model that estimated an increase in *Xist* expression with age (Model 1). We also looked for age-DE genes at the class level using only RFP+ neuron enriched libraries (thus removing any potential confounding of FACS population plan). We found that all neuronal subclasses have positive age effect sizes (**Extended Data Figure 4e**), supporting the ideal that the age effect size estimates from Model 1 are more accurate for the gene *Xist*. The reason we did not do this initially at the subclass level was due to lack of coverage of an adequate number of subclasses using only RFP+ libraries. As such, we chose to include libraries from many different FACS population plan collection strategies to maximize cell counts.

#### Adjusted age proportion calculation

We calculated the adjusted age proportion of each cluster by normalizing to the subclass-wide age proportion, as different brain regions profiled in this dataset vary in their proportions of aged versus adult cells (**Figure 2**). To do this, we subtracted the subclass-wide age proportion from the cluster-wide age proportion, and then added 0.5.

#### UMAP projection

We used principal components (PCs) calculated from PCA to calculate UMAPs for different groups of cells^91^. For UMAPs with >100,000 cells, we performed PCA based on the imputed gene expression matrix of genes based on top marker genes from each cluster within each grouping of cells as we have implemented previously^17^. For UMAPs with <100,000 cells, no imputation was used. Three parameters that can be adjusted when generating UMAPs include 1) number of PCs which are used to calculated projections, 2) nn.neighbors: the size of the local neighborhood of cells the UMAP will look at when trying to learn the structure of the data, 3) md: the minimum distance apart that cells are allowed to be in low dimensional resolution. For all UMAPs, the top 150 PCs were then selected, and PCs with >0.7 correlation were removed based on the technical bias vector, defined as log_2_(gene count) for each cell. Each PCA was run with unique gene list and each UMAP was run with a different set of nn.neighbors and md parameters. The parameters used for each PCA/UMAP are as follows: 6,446 genes, nn.neighbors = 10, md = 0.4 for the global UMAP (**Figure 1**); 984 genes, nn.neighbors = 20, md = 0.5 for the OPC-Oligo UMAP (**Figure 3**); 1,884 genes, nn.neighbors = 5, md = 0.5 for the Immune/Vasvular UMAP (**Figure 4**); 1,806 genes, nn.neighbors = 20, md = 0.5 for the Astro-Epen UMAP (**Figure 5**); 401 genes, nn.neighbors = 5, md = 0.5 for the tanycyte/ependymal cell UMAP (**Figure 5**); 1,169 genes, nn.neighbors = 5, md = 0.5 for the HY neuron UMAP (**Figure 6**).

#### Constellation plot

The global relatedness between cell types was visualized with constellation plots, which we had implemented previously^16,17^. To generate the constellation plot, each transcriptomic cluster was represented by a node (circle), whose surface area reflected the number of cells within the subclass in log_10_ scale. The position of each node was based on the centroid position of the corresponding cluster in UMAP coordinates. The relationships between nodes were indicated by edges that were calculated as follows. For each cell, 15 nearest neighbors in reduced dimension space were determined and summarized by cluster. For each cluster, we then calculated the fraction of nearest neighbors that were assigned to other clusters. The edges connected two nodes in which at least one of the nodes had > 5% of nearest neighbors in the connecting node. The width of the edge at the node reflected the fraction of nearest neighbors that were assigned to the connecting node and was scaled to node size. For all nodes in the plot, we then determined the maximum fraction of “outside” neighbors and set this as edge width = 100% of node width. The function for creating these plots, *plot_constellation* included in the R package scrattch.bigcat.

#### Gene ontology analysis

Gene ontology term enrichment was performed using the R package **clusterProfiler 4.0**^92^ and **gprofiler2**^93^. The function *gconvert* from gprofiler2 was used to convert gene IDs to their Ensmbl IDs. The functions *enrichGO* and *simplify* from clusterProfiler were then used to enrich for gene ontology terms from all three GO databases (molecular function, biological process, and cellular component). A p-value cutoff of 0.05 was used to determine significant GO terms.

### *In situ* spatial transcriptomics

#### Resolve Molecular Cartography overview

All *in situ* spatial RNA data shown here were generated by Resolve Biosciences with their commercially available Molecular Cartography platform. Two total Molecular Cartography experiments were performed (RSTE1-2), each with a different panel of 100 genes and targeting different region(s) of the brain (**Extended Data Figure 3**). For RSTE1, 4 different regions of the brain (cortex, striatum, midbrain, and hindbrain) were imaged in both sexes and both ages (2- and 18-month), with 2 replicate brains per condition and 2 technical replicates per brain. The technical replicates were plotted and analyzed as independent replicates in all figures. For RSTE2, the hypothalamus was imaged in both sexes and both ages, with 4 replicate brains per condition. Brain dissection and cryosectioning for Molecular Cartography experiments were performed at the Allen Institute for Brain Science in Seattle, WA, samples were stored at −80°C for 1-3 days, and then shipped overnight to Resolve Biosciences in San Jose, CA, where the Molecular Cartography protocol was performed. Spot data were then made available 1-2 weeks after receipt of tissue. Data analysis was performed at the Allen Institute using methods detailed below. Briefly, transcript data were segmented into cells, cells were filtered based on quality metrics generated from segmentation and mapping, and downstream analysis and visualization was performed.

#### Brain dissection and freezing

Mice used for spatial experiments were housed and kept in same conditions to those used for scRNA-seq described above. Mice were transferred from the vivarium to the procedure room with efforts to minimize stress during transfer. Mice were anesthetized with 5% isoflurane. A grid-lined freezing chamber was designed to allow for standardized placement of the brain within the block in order to minimize variation in sectioning plane. Chilled OCT was placed in the chamber, and a thin layer of OCT was frozen along the bottom by brief placement of the chamber in a dry ice/ethanol bath. The brain was rapidly dissected and placed into the prechilled OCT for approximately 2 minutes to acclimate to the cold prior to freezing. The orientation of the brain was adjusted under a dissecting scope, and the freezing chamber containing OCT and brains was placed into a dry ice/ethanol bath for freezing. After freezing, the brains were vacuum sealed and stored at −80°C.

#### Cryosectioning

The fresh-frozen adult and aged brains were sectioned at 10-µm on Leica 3050 S cryostats. The OCT block containing a fresh frozen brain was trimmed in the cryostat until reaching the desired region of interest. Sections were placed onto coverslips provided by Resolve Biosciences. Two replicate sections were collected sequentially – one as the primary sample and the other as a backup.

#### Gene panel design

The Molecular Cartography platform allows 100 genes per experiment for spatial RNA profiling. Each of the 2 Molecular Cartography experiments we ran was designed to target different regions and cell types in the adult and aged brains. Therefore, for each experiment we used different gene panels, which were compiled through a combination of automated and manual processes. Glutamatergic and GABAergic neuronal class markers *Slc17a7*, *Slc17a6*, *Gad1*, and *Gad2* and major non-neuronal subclass markers *Aqp4, Apod, Sox10, Pdgfra, Enpp6, Opalin, Dcn, Pecam1, Ctss, Mrc1, Kcnj8, Pdgfrb,* and *Acta2* were included for all 2 Resolve experiments. The remaining genes in each panel were then customized for each of the 2 experiments. RSTE1 targeted non-neuronal types in different parts of the brain. RSTE2 targeted tanycytes and ependymal cells in the third ventricle of the hypothalamus. The function *select_N_markers* included in the R package scrattch.hicat was used to select markers for all relevant subclasses and clusters in each experiment. Top age-DE genes were also included for relevant subclasses within each panel, as well as additional genes of interest selected from prior literature.

#### Cell segmentation

Cells were segmented using a combination of open source software **Cellpose**^94^ and **Baysor**^95^. Cellpose employs a generalist algorithm for segmenting cells from images of cellular stains as input. Baysor uses a transcript-driven algorithm to draw cell boundaries based on transcript data alone while also having the option of integrating prior knowledge from stained images into the process. First, images of DAPI stains from each of the tissue samples were used as input for Cellpose using the following parameters: *--pretrained_model = nuclei*, *--diameter = 0*. The output of Cellpose was saved as a TIF and used as a prior for the Baysor segmentation algorithm. Baysor was run with the following input parameters: *-m 30, -s 50*.

#### *In situ* data pre-processing

All segmented cells were mapped to the Allen whole mouse brain cell type taxonomy^17^ with the same method used for scRNA-seq data as described above. The 2 RSTE datasets were filtered for high-quality cells using a combination of thresholds for mapping confidence score, segmentation confidence score (from Baysor), number of transcripts, and gene detection. Due to the variable gene panels and brain regions across the two RSTE datasets, we used a different set of filter criteria for each experiment. These cutoffs are detailed in **Supplementary Table 6** and cell counts before and after quality filtering are diagramed in **Extended Data Figure 3**.

## Acknowledgments

We are grateful to the Transgenic Colony Management, Lab Animal Services, Molecular Biology, Tissue Processing, and Histology teams at the Allen Institute for technical support. We thank Dong-Wook Kim, John Mich, and JoAnn Buchanan for their feedback on the manuscript. The research was funded by grants from the National Institutes of Health (NIH), specifically, grant R01AG066027 from National Institute on Aging to H.Z. and B.T., and BRAIN Initiative grant U19MH114830 from National Institute of Mental Health to H.Z. The content is solely the responsibility of the authors and does not necessarily represent the official views of NIH and its subsidiary institutes. This work was also supported by the Allen Institute for Brain Science. The authors thank the Allen Institute founder, Paul G. Allen, for his vision, encouragement, and support.

## Author Contributions

Conceptualization: H.Z., B.T. Data generation and analysis lead: K.J. Data generation (scRNA-seq): K.J., Z.Y., C.T.J.vV., S.T.B., E.B., D.C., T.C., M.C., M. Departee, M. Desierto, J. Gloe, N.G., J. Guzman, D.H., E.L., T.P., M.R., K.R., J. Sevigny, N.S., L.S., J. Sulc, A.T., H.T., B.L., N.D., K.A.S., B.T., H.Z. Data processing and analysis (scRNA-seq): K.J., Z.Y., C.T.J.vV., A.B.C., R.C., J. Goldy, C.L., K.A.S., B.T., H.Z. Data generation (Resolve): K.J., A.G., A.R., B.T., H.Z. Data processing and analysis (Resolve): K.J., Z.Y., B.T., H.Z. Project management: E.S.K., K.G., S.M.S., K.A.S., L.E. Management and supervision: Z.Y., C.T.J.vV., B.L., S.M.S., N.D., L.E., K.A.S., B.T., H.Z. Manuscript writing and figure generation: K.J., Z.Y., C.T.J.vV., B.T., H.Z. Manuscript review and editing: K.J., Z.Y., C.T.J.vV., E.S.K., B.T., H.Z.

## Data Availability

Primary data will be deposited to the Neuroscience Multi-omic Data Archive (NeMO), https://nemoarchive.org/.

## Code Availability

Analysis methods used in the manuscript from R package **scrattch.hicat** and **scrattch.bigcat**, are available via github https://github.com/AllenInstitute/scrattch.bigcat.

---

**Extended Data Figure 1:**
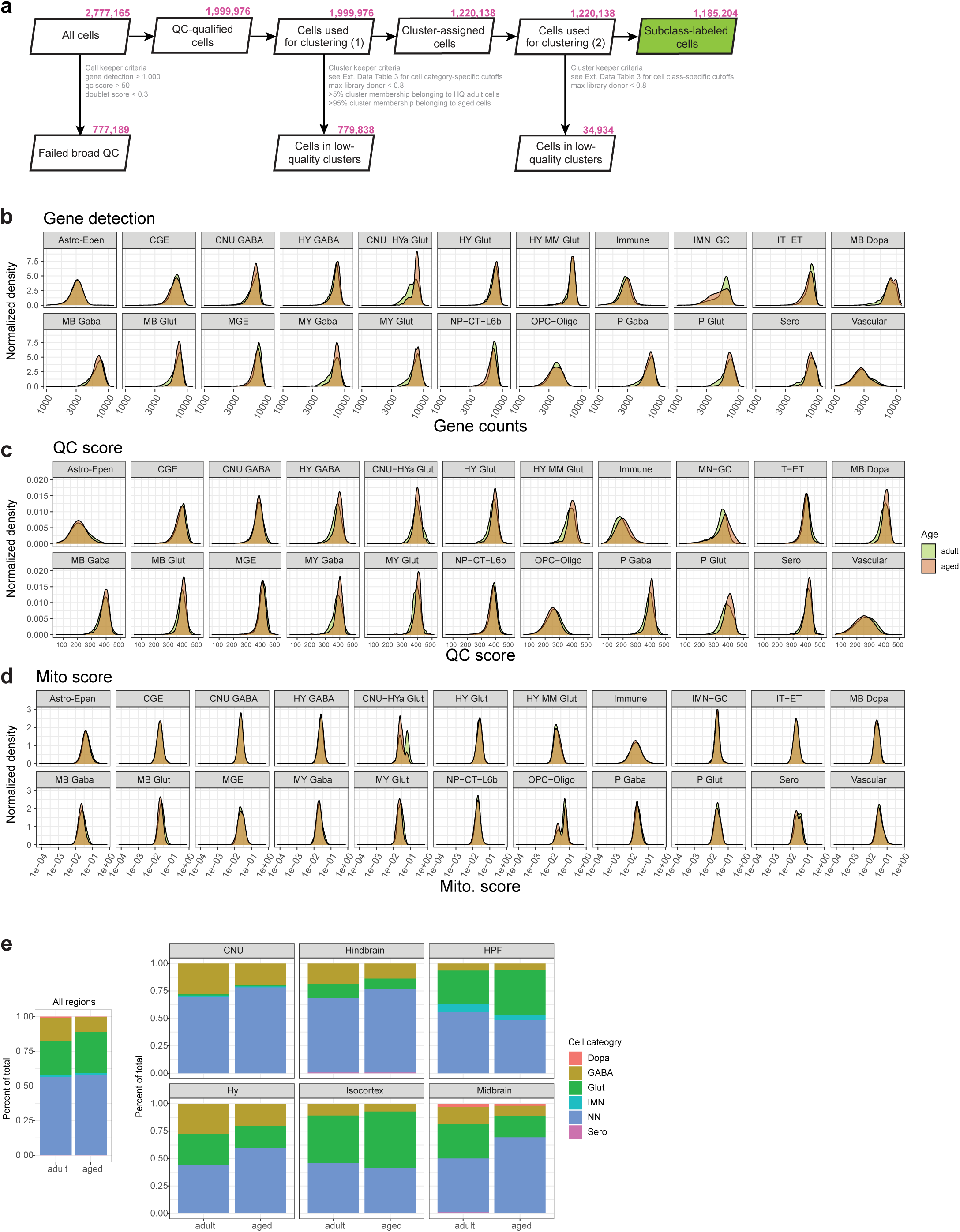
Data pre-processing workflow and quality control. **(a)** Workflow for pre-processing of scRNA-seq data. Cells retained at each step are indicated in pink. **(b-d)** Normalized density distribution of gene detection (b), QC score (c), and mito. score (d) per cell across different cell classes and ages. **(e)** Proportion of cell categories across all regions and within each major brain structure. Cell category: Dopa, dopaminergic neurons; GABA, GABAergic neurons; Glut, glutamatergic neurons; IMN, immature neurons; NN, non-neuronal cells; Sero, serotonergic neurons.

**Extended Data Figure 2:**
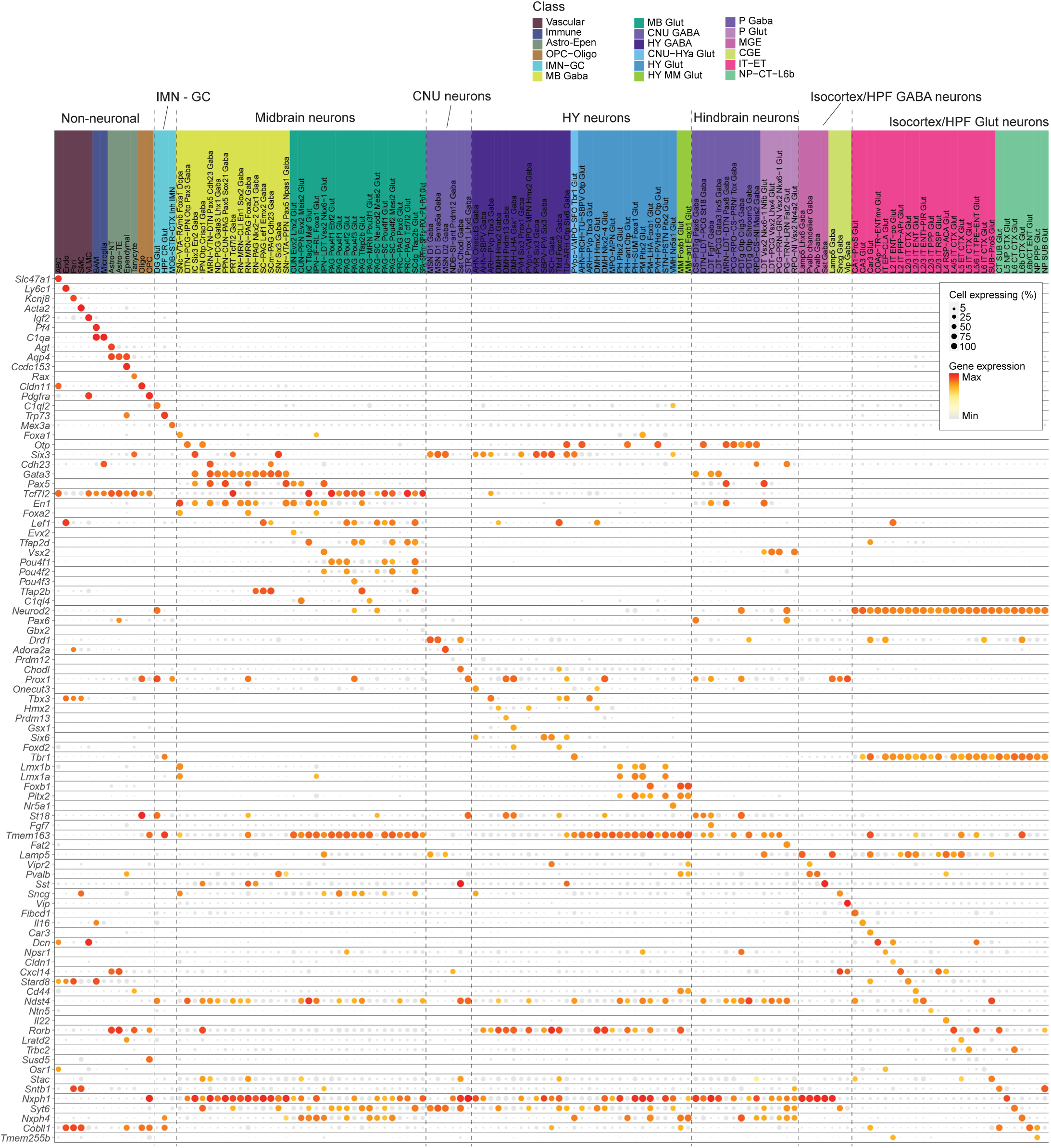
Subclass marker genes. Dot plot of marker gene expression for 132 individual subclasses of cell types analyzed in this study. Dot size and color indicate proportion of expressing cells and average expression level in each subclass, respectively. Subclass labels are colored by cell class.

**Extended Data Figure 3:**
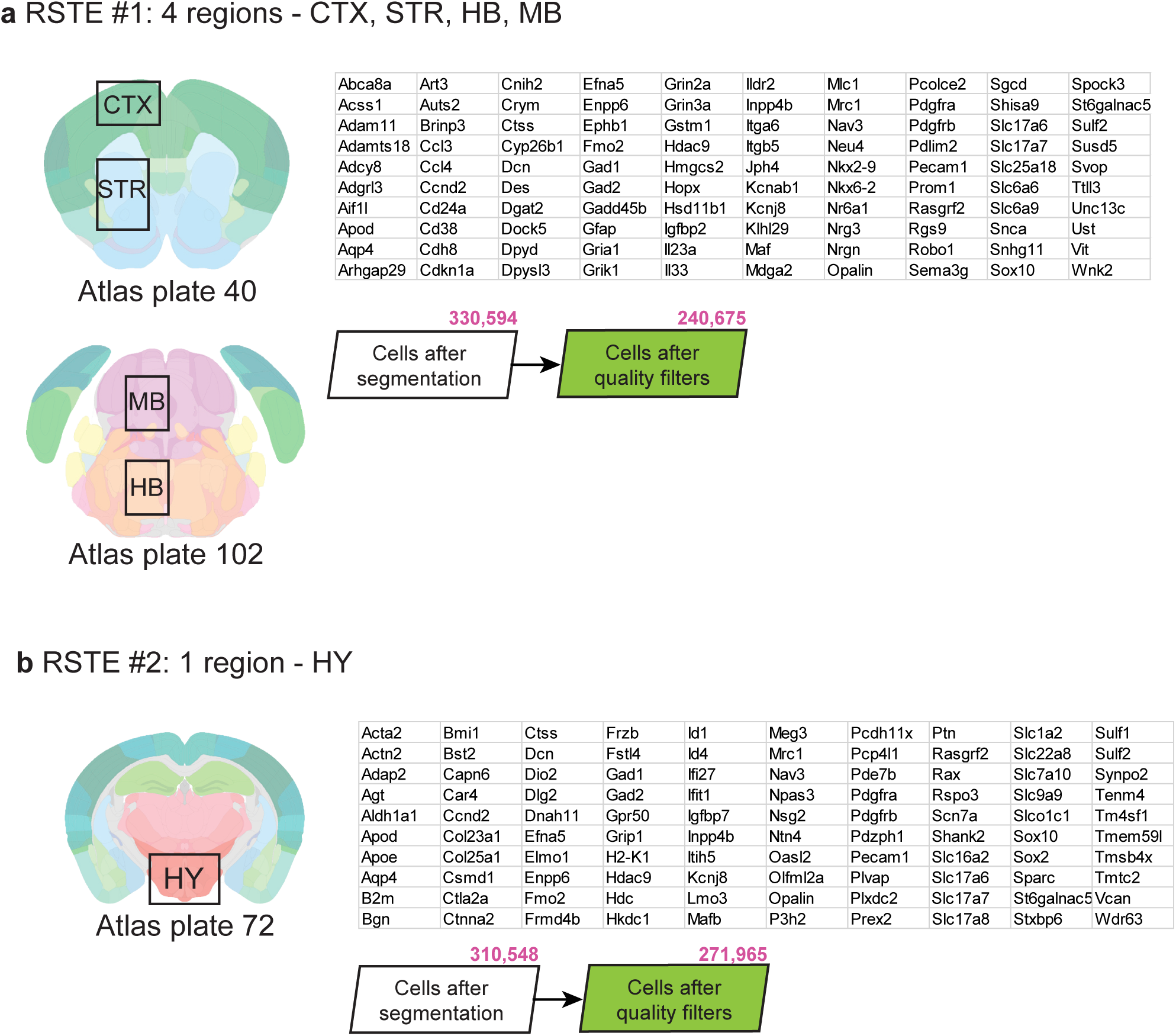
Summary of spatial transcriptomics datasets. **(a-b)** Diagram of brain regions profiled, gene panels, and pre- and post-filtered cell counts of Resolve spatial transcriptomic datasets 1 (RSTE1; a) and 2 (RSTE2; b).

**Extended Data Figure 4:**
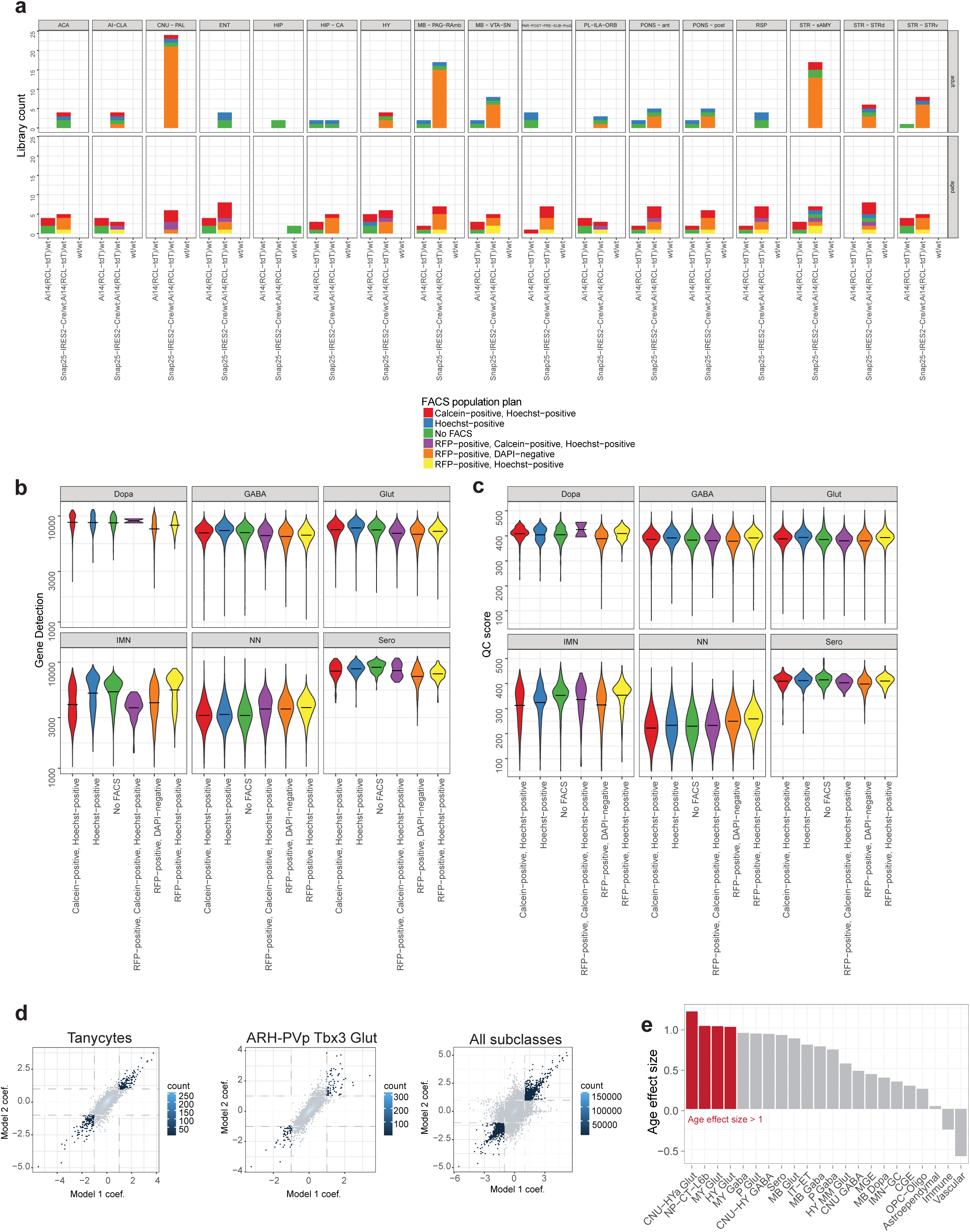
Library breakdown and DE gene model. **(a)** Summary of the numbers of libraries colored by FACS population plan and grouped by genotype (x-axis), age (rows), and ROI (columns). **(b-c)** Violin plot summary of gene detection (b) and QC score (c) grouped by FACS population plan (x-axis) and major cell category. **(d)** Two-dimensional density scatter plots of age effect sizes (coef) from simple and complex DE gene models plotted against one another for tanycytes only, ARH-PVp Tbx3 Glut neurons only, or all subclasses. Greater density is marked by lighter blue color. Dotted lines indicate significant cutoffs used in this study. Genes that pass these cutoffs are included in this study and summarized in Figure 2. (**e**) Bar plot of the age effect sizes of the gene *Xist* in decreasing order for all classes with n > 50 cells from each age and sex from RFP+, DAPI-libraries only. Significant changes (age effect size > 1 & p < 0.01) are colored in red.

**Extended Data Figure 5:**
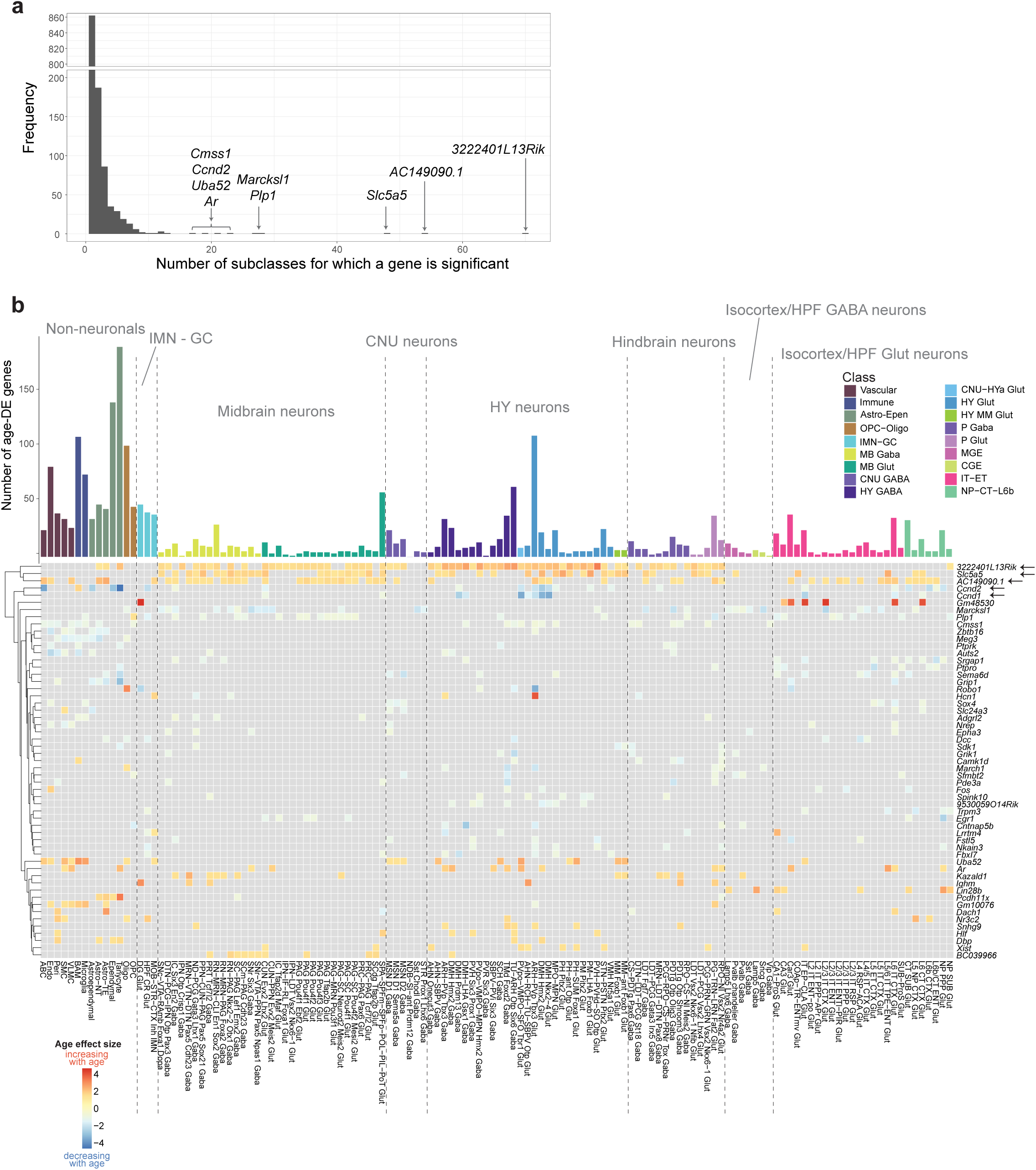
Common age-DE genes across subclasses. **(a)** Histogram of the number of subclasses an age DE gene is significant for. **(b)** Summary of the most commonly observed age-DE genes across all subclasses. Top: Summary of total age-DE genes colored and ordered by cell class, identical to that shown in Figure 2. Bottom: Heatmap of age effect sizes of the most common significant age-DE genes. DE genes that are significant in >5 subclasses are included. Genes are hierarchically clustered based on age effect size and their relatedness represented by the dendrogram.

**Extended Data Figure 6:**
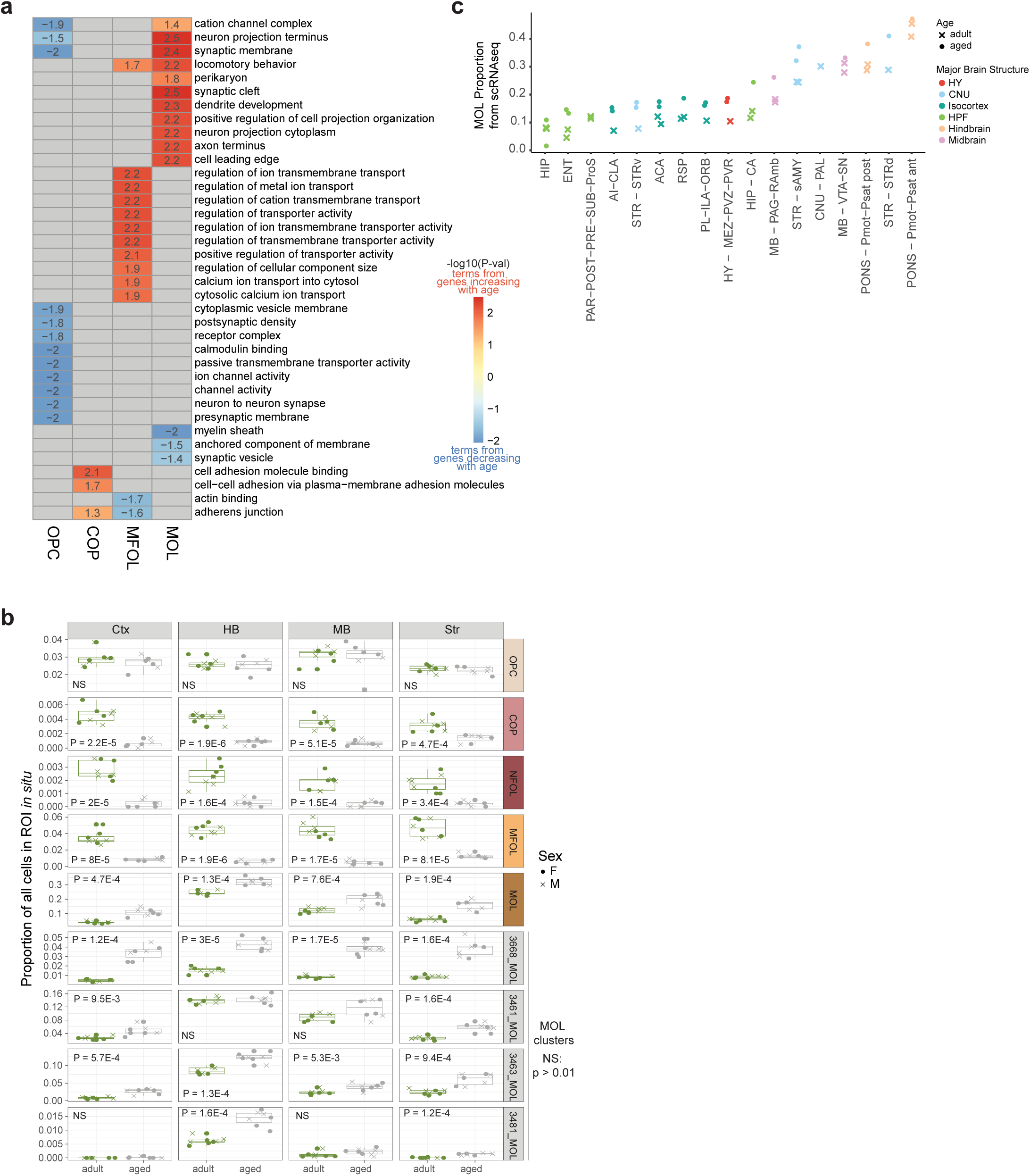
GO terms and changes in proportions in oligodendrocyte supertypes. **(a)** Heatmap of the statistical significance of top GO terms enriched in top age-DE genes from oligodendrocyte supertypes. Terms that are enriched in genes that increase with age are colored redder, while terms enriched in genes that decrease with age are colored bluer. Numbers in the plot represent -log10(p-value) of each term. **(b)** Relative changes in abundance of different supertypes and MOL clusters with age, calculated from spatial dataset RSTE1. A cutoff of p < 0.01 was used to determine statistical significance (Student’s t-test; NS, not significant). Each point corresponds to a replicate mouse sample. **(c)** Proportional changes of MOL with age, calculated from unbiased scRNA-seq libraries (i.e., libraries processed with the “No FACS” method). Each point represents one scRNA-seq library.

**Extended Data Figure 7:**
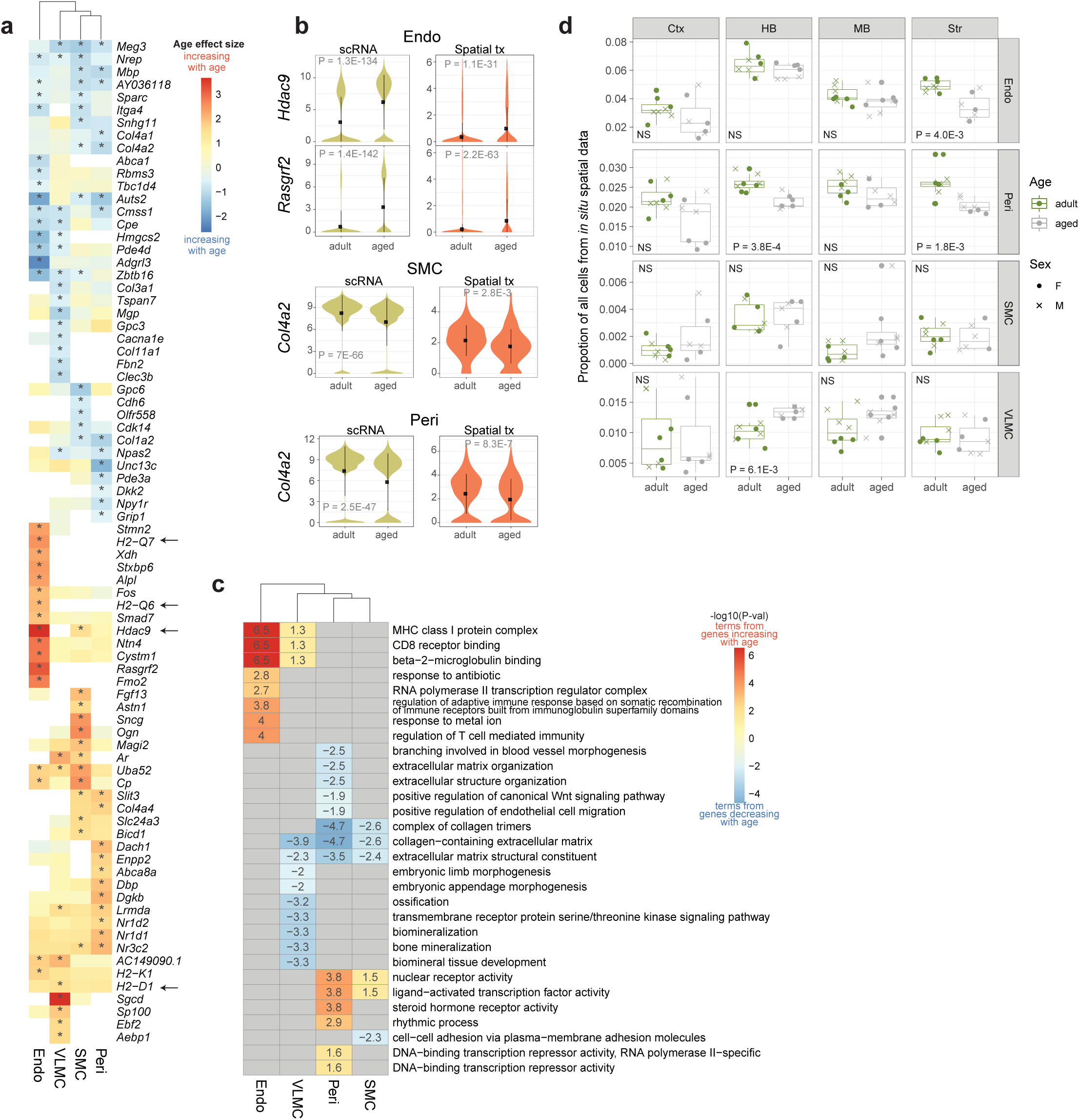
Age-associated changes in vascular types. **(a)** Heatmap of age effect sizes of top age-DE genes in Endo, VLMC, SMC, and Peri subclasses. Asterisk denotes statistical significance. Subclasses are hierarchically clustered based on age effect sizes and represented by the top dendrogram. **(b)** Violin plot expression of *Col4a2* in SMC and Peri subclasses, and *Hdac9* and *Rasgrf2* in Endo in scRNA-seq and spatial RSTE1 datasets. **(c)** Heatmap of the statistical significance of top GO terms enriched in top age-DE genes from vascular subclasses. Terms that are enriched in genes that increase with age are colored redder, while terms enriched in genes that decrease with age are colored bluer. Numbers in the plot represent -log10(p-value) of each term. Subclasses are hierarchically clustered based on scores and their relatedness represented by the dendrogram. **(d)** Proportional changes of vascular cell types with age calculated from spatial dataset RSTE1. Statistical significance is calculated with student’s t-test. Each point represents a single spatial replicate mouse sample.

**Extended Data Figure 8:**
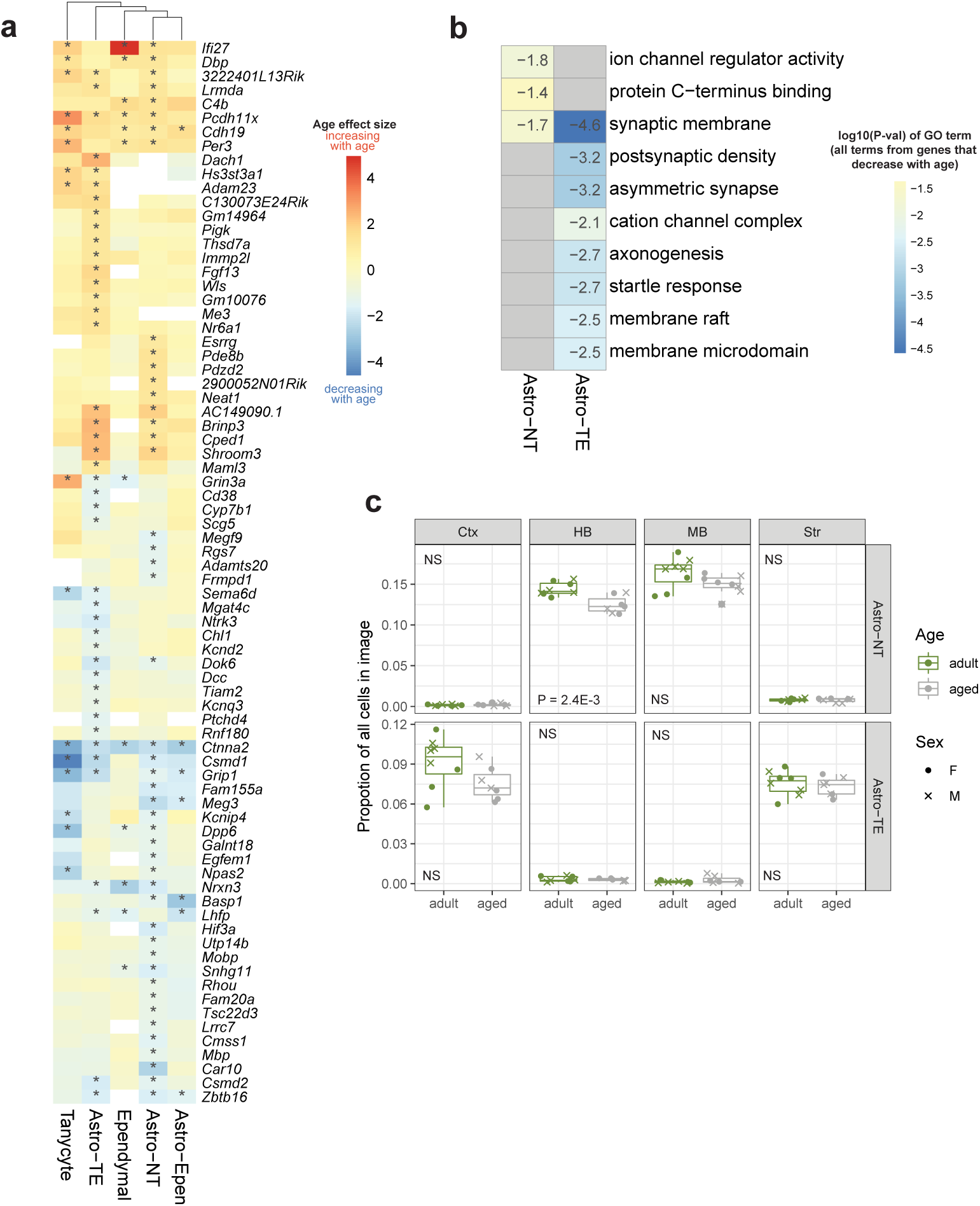
Age-associated changes in astrocytes. **a)** Heatmap of age effect sizes of top age-DE genes from Astro-TE and Astro-NT subclasses. Other Astro-Epen subclasses are included for reference. Asterisk denotes statistical significance. Subclasses are hierarchically clustered based on age effect sizes and represented by the top dendrogram. **(b)** Heatmap of the statistical significance of top GO terms enriched in top age-DE genes from Astro-TE and Astro-NT. All terms are enriched from genes that decrease with age. **(c)** Proportional changes of Astro-TE and Astro-NT cells with age calculated from spatial dataset RSTE1. Statistical significance is calculated with student’s t-test. Each point represents a single spatial replicate mouse sample.

## Supplementary tables

Supplementary Table 1: scRNA-seq library list

Supplementary Table 2: All cell subclasses analyzed in this study

Supplementary Table 3: All significant age-DE genes across subclasses, supertypes or clusters

Supplementary Table 4: All significant GO terms by sign at subclass, supertype or cluster levels

Supplementary Table 5: scRNA-seq cluster-level QC parameter cutoffs by cell type groupings

Supplementary Table 6: Resolve spatial transcriptomics cell-level QC parameter cutoffs by cell type groupings

